# Mitochondrial dynamics and mitophagy are necessary for proper invasive growth in Rice Blast

**DOI:** 10.1101/592915

**Authors:** Yanjun Kou, Yunlong He, Jiehua Qiu, Shu Yazhou, Fan Yang, YiZhen Deng, Naweed I. Naqvi

## Abstract

*Magnaporthe oryzae* causes Blast disease, which is one of the most devastating infections in rice and several important cereal crops. *M. oryzae* needs to coordinate gene regulation, morphological changes, nutrient acquisition, and host evasion, in order to invade and proliferate within the plant tissues. Thus far, the molecular mechanisms underlying the regulation of invasive growth *in planta* have remained largely unknown. We identified a precise filamentous-punctate-filamentous cycle in mitochondrial morphology during *Magnaporthe*-Rice interaction. Interestingly, loss of either the mitochondrial fusion (MoFzo1) or fission (MoDnm1) machinery, or inhibition of mitochondrial fission using Mdivi-1 caused significant reduction in *M. oryzae* pathogenicity. Furthermore, exogenous carbon source(s) but not antioxidant treatment delayed such mitochondrial dynamics/transition during invasive growth. Such nutrient-based regulation of organellar dynamics preceded MoAtg24-mediated mitophagy, which was found to be essential for proper biotrophic development and invasive growth *in planta*. We propose that precise mitochondrial dynamics and mitophagy occur during the transition from biotrophy to necrotrophy, and are required for proper induction and establishment of the blast disease in rice.

## INTRODUCTION

Mitochondria, the semi-autonomous double-membrane bound organelles, generate most of the adenosine triphosphate (ATP) for diverse cellular functions and are involved in various physiological processes including lipid metabolism, redox signalling, calcium- and iron-homeostasis, and programmed cell death (Zemirli & Morel, 2018, Nunnari & Suomalainen, 2012). Depending on the cellular physiology and environment, mitochondria exhibit a variety of morphologies, ranging from elongated and interconnected networks to small spherical organelles. The mitochondrial shape is dynamic and depends on the balance between two opposing processes, fusion and fission, which occur continuously during the growth cycle (Westermann, 2010). Maintaining the mitochondrial morphology in steady state by the balance of fusion and fission activities is critical for living cells. When this equilibrium is broken, mitochondrial shape and dynamics are disturbed leading to important physiological consequences including increased cellular stress and various diseases (Zemirli & Morel, 2018, Rapaport *et al.*, 1998, Guan *et al.*, 1993, Sesaki & Jensen, 2001, Chang & Doering, 2018, Ma *et al.*, 2009, Mozdy *et al.*, 2000, Delettre *et al.*, 2000, Kijima *et al.*, 2005).

The fusion and fission machineries of mitochondria are well conserved from yeast to mammals. In yeast, mitochondrial fusion mainly depends on the transmembrane GTPase Fzo1, membrane anchored dynamin GTPase Mgm1, and Ugo1, which links the outer and inner membrane fusion machineries (Rapaport et al., 1998, Guan et al., 1993, Sesaki & Jensen, 2001). In yeast, the loss of Fzo1, Mgm1, or Ugo1 leads to numerous small fragmented mitochondria due to a block in fusion and amidst ongoing fission of mitochondria (Rapaport et al., 1998, Guan et al., 1993, Sesaki & Jensen, 2001). The Fis1-Mdv1/Caf4-Dnm1 complex constitutes the major mitochondrial fission pathway in yeast (Mozdy et al., 2000, Griffin *et al.*, 2005). Fis1, a tail-anchored outer membrane protein, functions as a membrane receptor, and Mdv1/Caf4 serves as an adaptor to recruit dynamin-related protein Dnm1 to the fission sites in mitochondria (Mozdy et al., 2000, Griffin et al., 2005). Dnm1 is the key mediator of membrane scission during mitochondrial division (Mozdy et al., 2000). Loss of either Fis1 or Dnm1 blocks fission resulting in highly interconnected fishnet-like mitochondria (Mozdy et al., 2000).

In addition to fusion and fission machineries, mitochondrial homeostasis requires proper mitophagy, which is the selective sequestration of mitochondria by autophagosomes followed by their degradation in vacuoles/lysosomes (Liu *et al.*, 2014). Mitophagy is a key mechanism in organellar quality control, and is responsible for the removal of damaged or unwanted mitochondria (Liu et al., 2014). In yeast, the mitochondrial outer membrane receptor Atg32 is essential for mitophagy (Kanki *et al.*, 2009, Okamoto *et al.*, 2009). In response to nitrogen starvation or inhibition of mTOR following growth in non-fermentable carbon source, Atg32 directs mitochondria to the autophagosome through its interaction with the core autophagic machinery, including Atg8 and Atg11, to induce mitophagy (Kanki et al., 2009, Okamoto et al., 2009). Such receptors sense stimuli that induce mitophagy, and couple mitochondrial dynamics to the quality control machinery (Mao & Klionsky, 2013).

Although the fusion and fission machineries are highly conserved, diverse mechanisms ensure proper organellar dynamics and distribution to optimize mitochondrial function in response to changing environments and cellular needs. The entire mitochondrial network is fused during G1-S phase transition and fragmented/punctate in the late S and M phase depending on cellular environment in rat kidney cells (Mitra *et al.*, 2009). In addition, the mitochondrial dynamics are modulated in response to certain types and/or severity of stresses and adapt their form by promoting fusion or fission (Shutt & McBride, 2013). When cells are subjected to mild stresses, such as moderate nutrient starvation, protein synthesis inhibition, or mTOR inhibition induced autophagy, mitochondria tend to become more fused to increase ATP production and escape from mitophagy (Tondera *et al.*, 2009, Gomes *et al.*, 2011, Li *et* al., 2015). Conversely, the mitochondrial fission machinery is activated upon prolonged nutrient stress, leading to degradation via mitophagy or apoptosis (Toyama *et al.*, 2016, Frank *et al.*, 2001). It is clear that mitochondrial dynamics and mitophagy are directly associated with metabolic status and stress conditions (Twig *et al.*, 2008, Mao & Klionsky, 2013, Toyama et al., 2016).

*M. oryzae* is a hemibiotroph, which initially establishes a close biotrophic association to acquire nutrients from the live host cells, but later on switches to the necrotrophic killing phase to obtain nutrients from dead plant tissues (Fernandez & Orth, 2018). During the infection cycle in *M. oryzae*, the three-celled conidia are deposited by rain splashes and stick to the rice leaf surface. Under proper conditions, such conidia germinate and form appressoria to assist in the breach of the rigid rice cuticle. Once inside the host cell, *M. oryzae* differentiates into invasive hyphae and spreads to neighbouring cells resulting in typical lesion formation. During invasive growth, *M. oryzae* needs to coordinate the nutrient sensing, gene expression regulation, morphological changes, acquiring nutrients from rice cells, and eluding the plant immunity to adapt to the host milieu (Marroquin-Guzman *et al.*, 2017). The molecular mechanisms involved in regulating mitochondrial homeostasis during invasive growth have not been explored in depth. Recent studies have shown that the complex composed of MoDnm1, MoFis1, and MoMdv1 regulates the mitochondrial fission in *M. oryzae* (Zhong *et al.*, 2016). Disruption of *MoDNM1* or *MoFIS1* results in defects in mitochondria fission and pathogenicity (Zhong et al., 2016, Khan *et al.*, 2015). Our recent analyses showed that the sorting nexin MoAtg24 is essential for mitophagy and necessary for proper asexual differentiation (He *et al.*, 2013). However, the regulation and function of mitochondrial dynamics and mitophagy during *M. oryzae* development *in planta* need to be further explored.

In this study, a unique filamentous-punctate-filamentous cycle in mitochondrial morphology and dynamics was observed during the early infectious growth of *M. oryzae*. To uncover the role of this specific cycle and mitophagy, mutants defective in mitochondrial fusion, fission, and mitophagy were generated via deletion of *MoFZO1, MoDNM1,* or *MoATG24* respectively. Characterization of *Modnm1*Δ, *Mofzo1*Δ, and *Moatg24*Δ strains and Mdivi-1-based inhibition of mitochondrial division revealed that mitochondrial fusion and fission machineries and mitophagy are required for maintaining mitochondrial dynamics, and are necessary for proper infection and pathogenesis in *M. oryzae*. We provide evidence that carbon source depletion triggers such specific mitochondrial dynamics during the early infection stage. Overall, our study demonstrates that tightly controlled mitochondrial dynamics and mitophagy are required for proper invasive growth during establishment of the blast disease in rice.

## RESULTS

### Mitochondrial dynamics during *M. oryzae*-rice interaction

We first examined mitochondrial morphology during *in planta* growth of wild-type (WT) *M. oryzae* using the *Mito-GFP* (as the mitochondrial marker (He et al., 2013, Patkar *et al.*, 2012)) strain. The conidia of the *Mito-GFP* strain were inoculated on sheath from 21 d old susceptible rice seedling (*Oryza sativa* L., cultivar CO39), and incubated in a humid chamber at room temperature. Mitochondrial morphology was examined at the following three time points post inoculation: 30 hpi when the fungus successfully penetrated the rice epidermis, 48 hpi when most of invasive hyphae spread into the neighbouring rice cells and necrotrophy starts to occur, and 72 hpi when necrotrophy/lesion formation could be observed. At 30 hpi, the majority of mitochondria (81.2% ± 1.9%) were in a tubular or filamentous network (Figure 1). In contrast, most mitochondria (83.9% ± 9.9%) were fragmented or punctate at 48 hpi (Figure 1). Interestingly, about half of the mitochondria (50.5% ± 5.9%) appeared to be filamentous or tubular again at 72 hpi (Figure 1). However, such specific and dynamic changes in mitochondrial network were not evident during appressorium formation (Figure S1). Such temporal and dramatic changes in mitochondrial morphology indicated that *M. oryzae* likely faces dynamic environmental or cellular changes that significantly impact mitochondrial form/function during the first 72 h of *in planta* growth.

**Figure 1.**
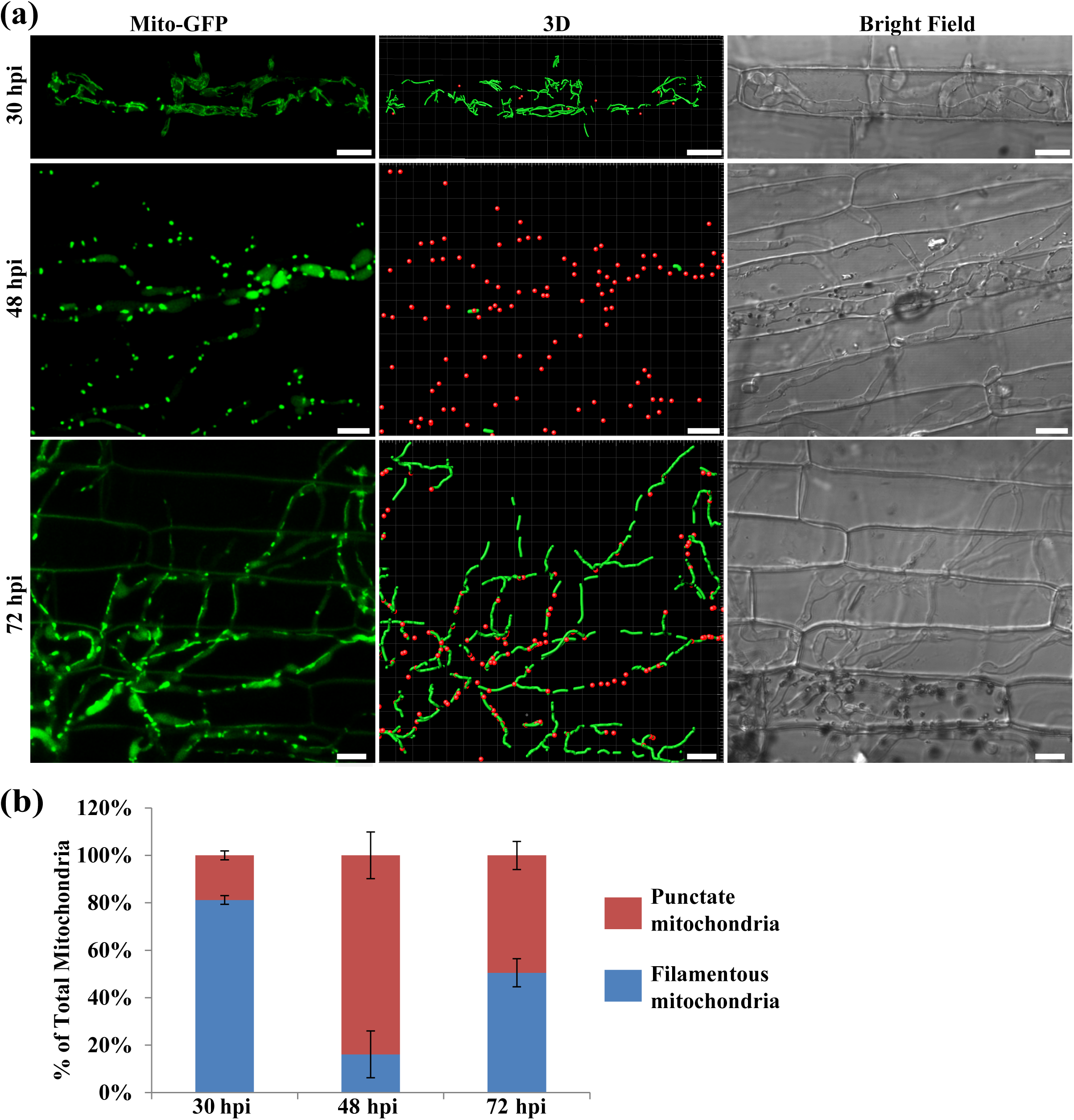
Specific changes in mitochondrial morphology during *in planta* growth of *M. oryzae*. (a) Mitochondrial morphology in *M. oryzae* during infection. The conidial suspension of the *Mito-GFP* strain was inoculated on rice sheath (*Oryza sativa* L., cultivar CO39). Confocal microscopy was carried at 30, 48, and 72 hpi. The 3D reconstruction of the mitochondrial morphology was performed in Bitplan Imaris. Red spots and green filaments represent punctate and filamentous mitochondria respectively. Scale bar: 8 μm. (b) Quantification of the different morphologies of mitochondria in the wild type *Mito-GFP* strain during infection. Error bars represent Mean ± SD from three independent replicates. Sample size is more than 200 appressoria penetration sites/host tissue per analysis.

### The role of mitochondrial dynamics in invasive growth in ***M. oryzae***

Mitochondrial dynamics through fusion and fission during invasive growth occurred prior to lesion development, raising the possibility that such organellar dynamics might play an important role in the establishment and spread of the blast disease. To determine the role of such changes in morphology and dynamics of mitochondria during *M. oryzae* infection, we generated mutants defective in mitochondrial fission (*Modnm1*Δ, Figure S2) or inhibited mitochondrial fission using Mdivi-1, or disrupted the mitochondrial fusion (*Mofzo1*Δ, Figure S3) to alter the overall mitochondrial network dynamics, and examined their invasive growth and the pathogenicity.

*MoDnm1* is known as an important mitochondria fission gene in *M. oryzae* (Zhong *et al*., 2016). In our study, *MoDnm1* was simply used as a marker gene for analyzing the loss of mitochondrial fission in *M. oryzae*. A gene-deletion mutant of *MoDNM1* was generated in the *Mito-GFP* strain. As previously reported (Zhong *et al*., 2016), the *Modnm1*Δ strain exhibited the characteristic tubular or fishnet-like mitochondrial structures (Figure 2a), suggesting that the *Modnm1*Δ is indeed incapable of mitochondrial fission. To verify the role of mitochondrial fission in fungal pathogenicity, the conidial suspension from WT, *Modnm1*Δ, or *Modnm1*Δ complemented strain was used for blast infection assays on rice seedlings. The *Modnm1*Δ strain showed highly reduced pathogenicity and formed small and highly restricted lesions at 7 dpi (Figure 2b, c). Furthermore, mitochondria in *Modnm1*Δ were tubular or filamentous at 30 hpi, 48 hpi, and 72 hpi, while a majority of mitochondria were fragmented/punctate in the WT at 48 hpi (Figure 3). Since the *Modnm1*Δ strain has pleiotropic defects in *M. oryzae*, we also performed the mitochondrial fission inhibitor treatment to confirm the role of mitochondrial fission during blast infection. Treatment with Mdivi-1, which inhibits mitochondrial fission in *M. oryzae* (Zhong et al., 2016), resulted in extensive tubular mitochondrial structures in *M. oryzae*, and significantly reduced the invasive growth in rice cells (Figure 4). Based on these results, we conclude that mitochondrial fission plays an important role in the invasive growth and lesion formation during Rice Blast.

**Figure 2.**
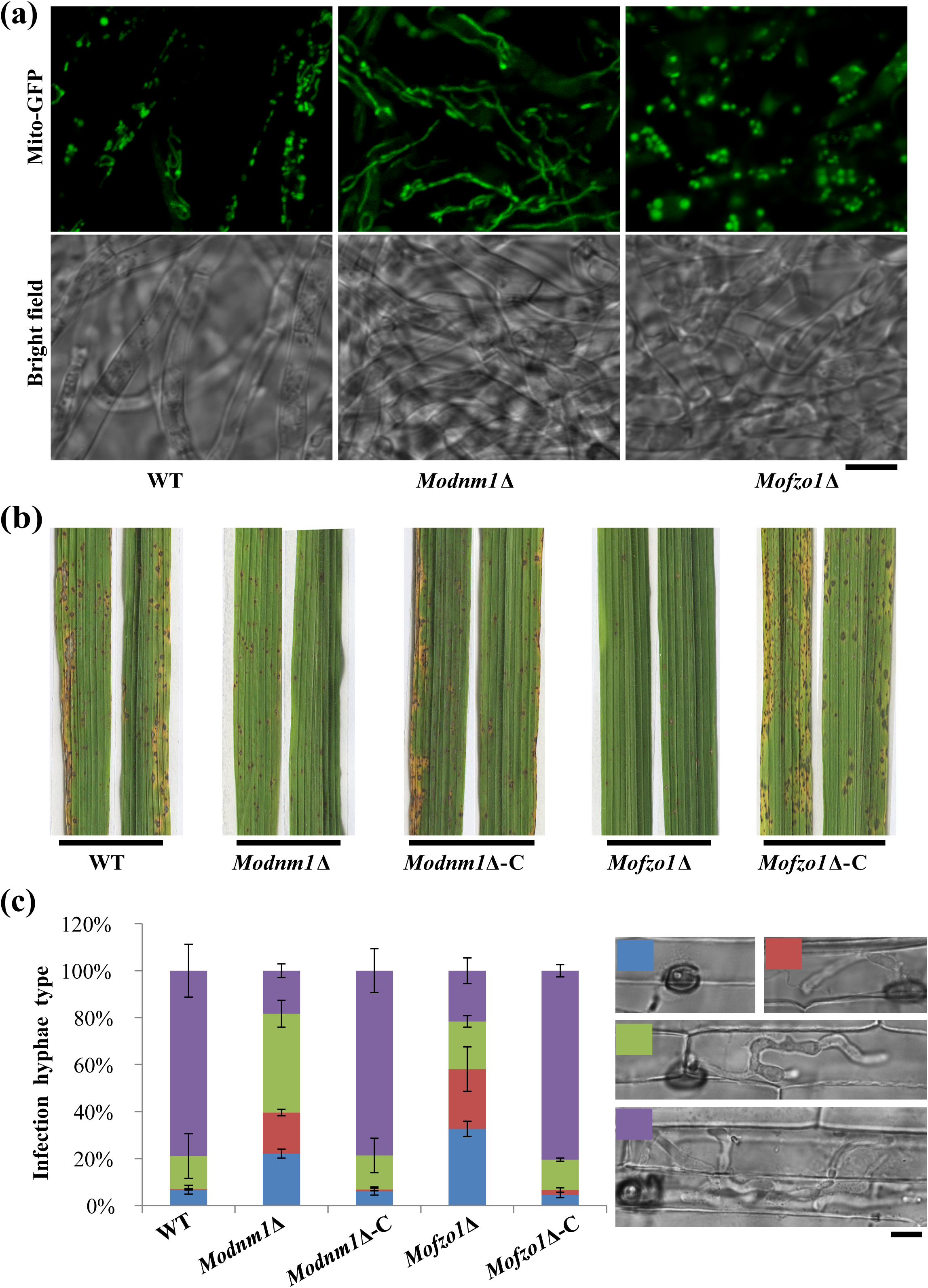
Mitochondrial fusion and fission are required for proper pathogenesis of *M. oryzae*. (a) The function of *MoDNM1* and *MoFZO1* in mitochondrial fission and fusion. Two day old liquid CM-grown mycelia of the indicated strains were used for imaging with confocal microscopy. Most of the mitochondria in *Modnm1*Δ formed elongated or interconnected fishnet-like structures, while the mitochondria were punctate or fragmented in *Mofzo1*Δ in vegetative mycelia. (b) The rice seedling (*Oryza sativa* L., cultivar CO39) infection assay of wild type (WT), *Modnm1*Δ, *Modnm1*Δ complementation strain (*Modnm1*Δ-C), *Mofzo1*Δ, and *Mofzo1*Δ complemented strain (*Mofzo1*Δ-C). (c) Detailed observation and statistical analysis of invasive growth in rice sheath cells at 40 hpi. Four types (illustrated in the right panel with corresponding colour labels): no penetration, penetration with primary hyphae, with differentiated secondary invasive hyphae, and invasive hyphae spreading into neighbouring cells, were quantified. Data represent mean ± SD of three independent experiments, with n = 200 appressoria per analysis. Scale bar represents 5 μm.

**Figure 3.**
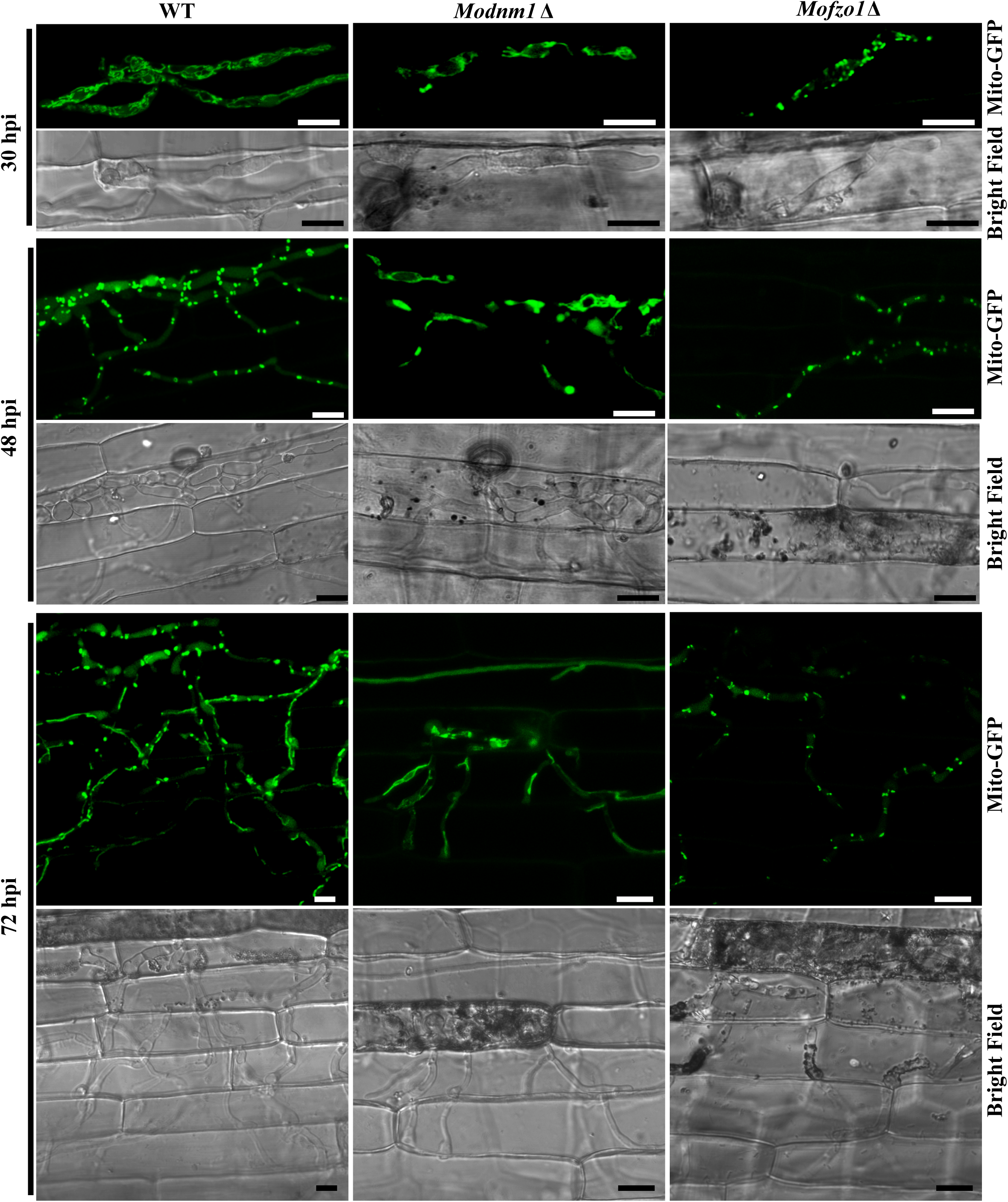
The mitochondrial morphology in WT, *Modnm1*Δ, and *Mofzo1*Δ during the infection process. Invasive hyphal growth of *Modnm1*Δ or *Mofzo1*Δ was significantly slower than WT. Scale bar = 10 μm.

**Figure 4.**
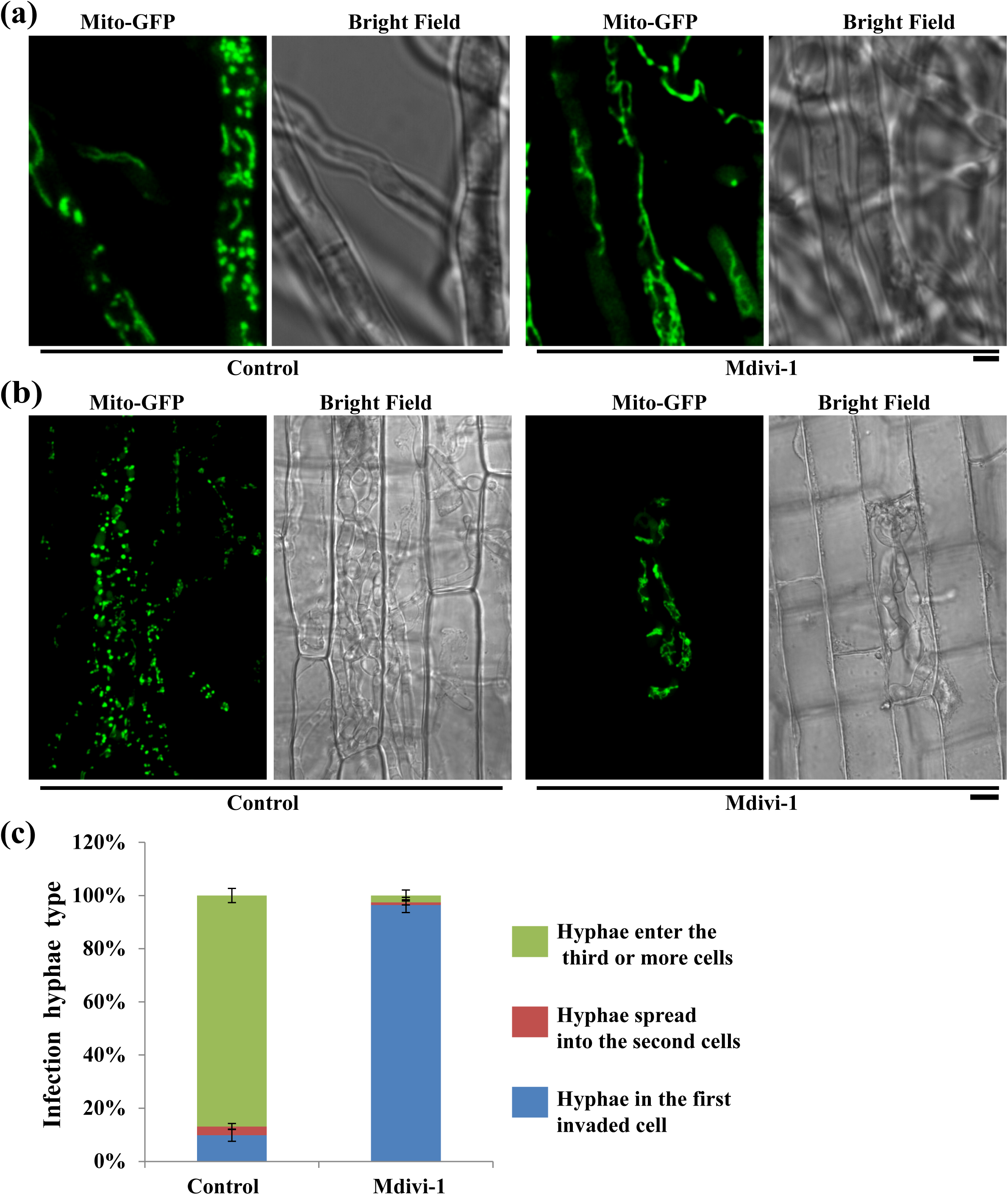
Chemical inhibition of mitochondrial fission reduces invasive hyphal growth of *M. oryzae*. (a) The mitochondria were predominantly tubular or filamentous upon Midvi-1 treatment. Scale bar represents 2 μm. (b) The mitochondrial morphology of the *Mito-GFP* strain with/without Midvi-1 at 48 hpi. Scale bar = 8 μm. (c) Detailed observation and statistical analysis of invasive growth in rice sheath cells at 48 hpi.

In *S. cerevisiae,* Fzo1 is the first known mediator of mitochondrial fusion (Rapaport et al., 1998, Fritz *et al.*, 2001). Deletion mutant of the orthologous *MoFZO1* harboured punctate mitochondria (Figure 2a), thus indicating a mitochondrial fusion defect in this *M. oryzae* mutant. Similar to *Modnm1*Δ, the *Mofzo1*Δ strain formed small and restricted blast lesions on rice plants (Figure 2b). Further microscopic observations showed that more than 90% of appressoria penetrated successfully and about 80% infectious hyphae extended to neighbouring cells in the WT and the complemented strain at 40 dpi, while only 67.5% of appressoria penetrated successfully and 21.6% invasive hyphae spread to surrounding cells in the *Mofzo1*Δ strain (Figure 2c; P<0.005). We further analyzed the mitochondrial dynamics during the blast infection process. As shown in Figure 3, the mitochondria in *Mofzo1*Δ were punctate at all the time points tested. These results indicated that mitochondrial fusion within the blast pathogen is required for proper invasive growth and lesion formation.

Taken together, we conclude that the mitochondrial fission and fusion machineries are involved in invasive growth in *M. oryzae*; and mitochondrial dynamics plays a crucial role during *Magnaporthe* pathogenesis.

### Carbon source depletion triggers mitochondrial fragmentation

Mitochondrial fragmentation can be triggered by multiple environmental factors such as oxidative stress, and carbon source depletion (Zemirli & Morel, 2018). During host invasion, the fungal pathogen generally encounters the plant defense response, oxidative stress, and metabolic stress. We therefore hypothesized that such host response, oxidative, and/or metabolic stress, triggers the specific mitochondrial fragmentation during invasive growth *in planta*.

To test whether the live host factors trigger such changes in mitochondrial fragmentation, we first examined the mitochondrial morphology in the blast fungus in live host tissue and compared it to that in heat-killed rice sheath. The heat treatment was used to first kill the rice sheath cells before inoculating with the blast fungal strain of interest. Mitochondrial fragmentation was evident in invasive hyphae in heat-killed rice sheath at 48 hpi. However, the percentage of filamentous mitochondria was significantly higher than the control samples at 48 hpi and did not show any difference at 72 hpi (Figure 5). These results indicated that the mitochondrial fragmentation observed during *M. oryzae* invasive growth is dependent in part on the active defense response in addition to other factors in live host plants.

**Figure 5.**
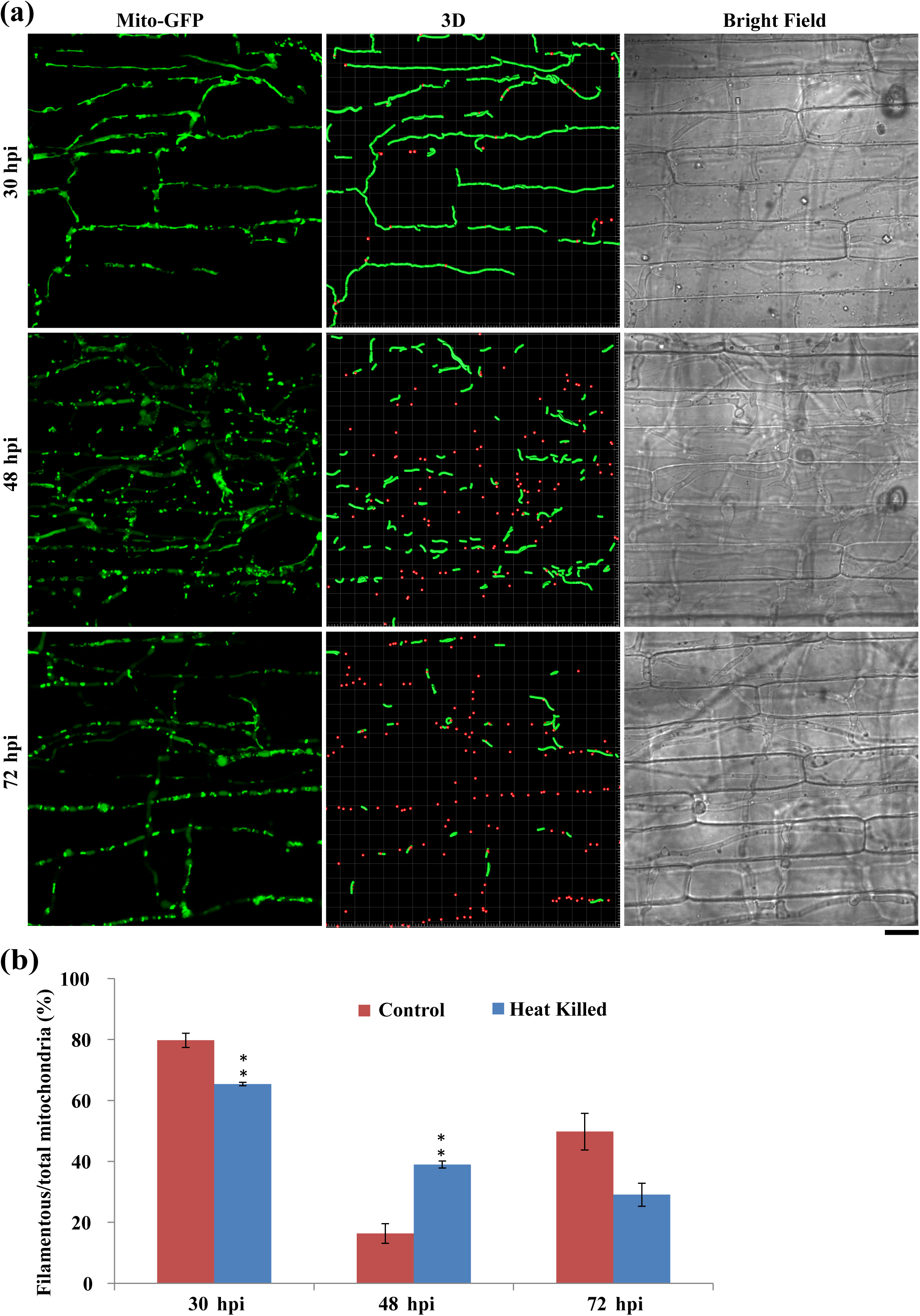
The mitochondrial morphology and dynamics in the *Mito-GFP* strain in heat-killed rice sheath. (a) Confocal microscopy images of the *Mito-GFP* strain in dead rice sheath cells at 30, 48, and 72 hpi. Scale bar = 12 μm. In the 3D image, red spots and green filaments highlight punctate and filamentous mitochondria respectively. (b) Quantitative analysis of mitochondria of different morphologies in the *Mito-GFP* strain in heat-killed rice cells. Values represent the mean ± SD from three independent experiments. Sample size is more than 200 appressoria penetration sites per analysis. **, P<0.005 compared with WT at indicated time point.

Oxidative stress could trigger the mitochondrial fragmentation in *M. oryzae* (Figure S4). We further tested whether oxidative stress triggers the mitochondrial fragmentation during invasive growth, by imaging the mitochondrial morphology at 30 hpi, 48 hpi, and 72 hpi in the presence of the exogenous antioxidant. The antioxidant treatment was initiated at 24 hpi. In the presence of 2.5 mM GSH (Glutathione) as an exogenous antioxidant, around 81% mitochondria still became punctate or fragmented at 48 hpi (Figure S5). At 72 hpi, filamentous or tubular mitochondria were apparent in GSH-treated invasive hyphae. Likewise, N-acetyl cysteine (NAC) treatment did not alter the mitochondrial fragmentation regime at 48 hpi (Figure S5). Therefore, we inferred that ROS/oxidative stress is unlikely to be the trigger for mitochondrial fragmentation during *M. oryzae* invasive growth.

Next, we examined the role of carbon source depletion on mitochondrial fragmentation as exogenous carbon sources have been reported to alter metabolic stresses (Toyama et al., 2016). The carbon source, glucose or sucrose, was individually added into inoculated conidia droplets on the rice sheath surface at 24 hpi. Mitochondrial morphology was assessed using a confocal microscope at 30 hpi, 48 hpi, and 72 hpi. We found that excess glucose or sucrose significantly delayed the mitochondrial fragmentation (Figure 6, 7), which indicated that carbon source depletion might be the major factor triggering mitochondrial fragmentation during *in planta* growth in *M. oryzae*. In the control experiments (no additional carbon source), around 84% mitochondria appeared fragmented at 48 hpi (Figure 6, 7), whereas mitochondrial fragmentation occurred at 72 hpi in the presence of the indicated exogenous carbon source. At 72 hpi, 69% and 67% of mitochondria were punctate upon additional supply of glucose or sucrose, respectively (Figure 6, 7), while more than 50% mitochondria in the WT appeared filamentous again. Based on these data, we conclude that the presence of excess carbon source impacts mitochondrial dynamics (and/or function) in invasive hyphae during blast development. Since the aforementioned carbon sources delay mitochondrial fragmentation to some extent, it is possible that downstream molecules in the carbon metabolic pathway regulate mitochondrial dynamics during blast infection. Accordingly, the important carbon metabolic intermediate G6P (Glucose-6-phosphate) was added to the inoculated conidial suspension at 24 hpi, and the mitochondrial morphology was assessed at 48 hpi and 72 hpi. Nearly 85% and 64% of total mitochondria remained tubular or filamentous in the presence of G6P around 48 hpi (P<0.001) and 72 hpi (Figure 6, 7; P<0.01). Taken together, these results indicate that carbon source depletion and the live host factors but not oxidative stress *per se*, trigger mitochondrial fragmentation during the invasive growth phase in *M. oryzae*.

**Figure 6.**
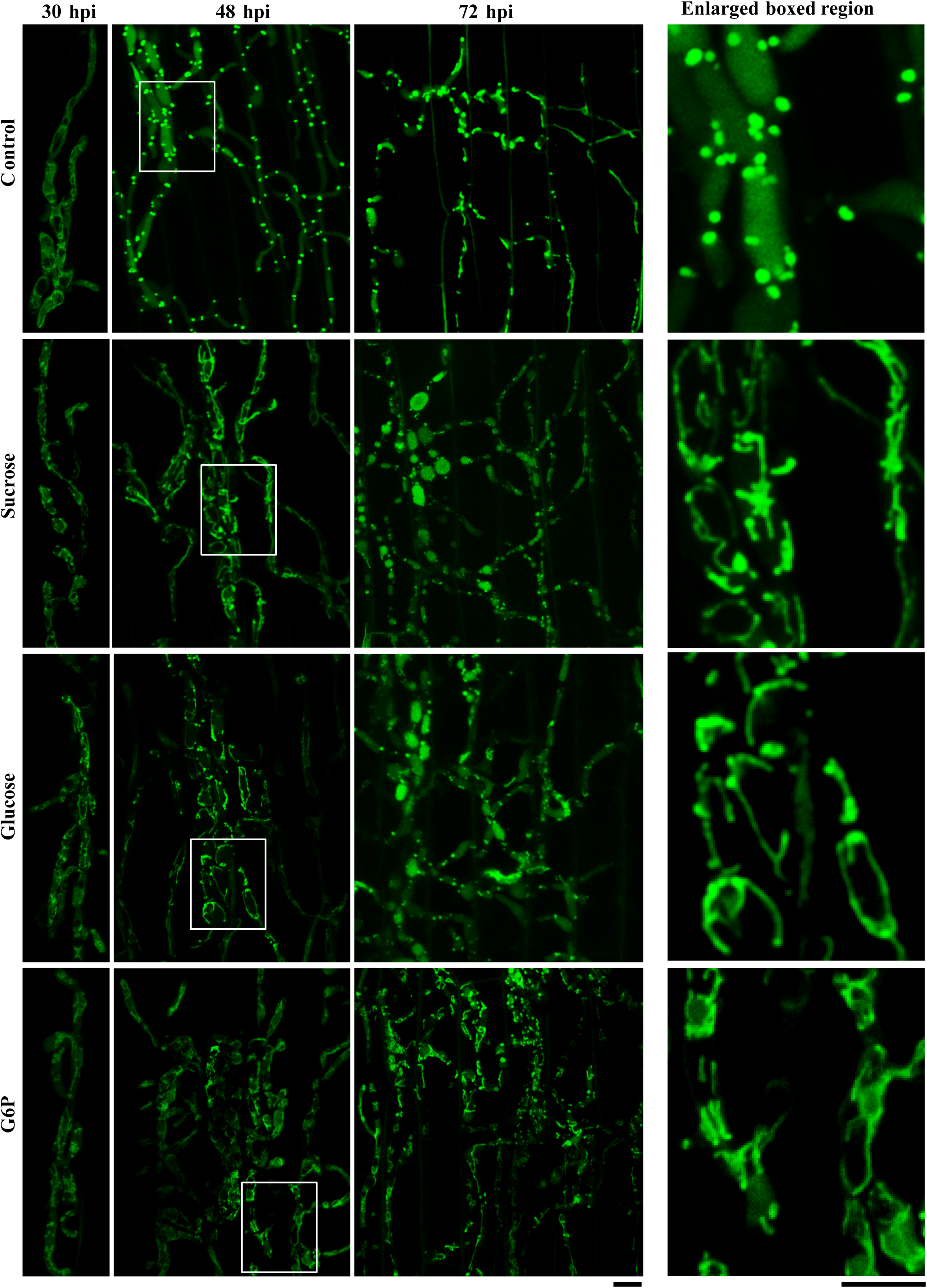
Carbon-replete condition delays mitochondrial fragmentation *in planta.* Conidia of the *Mito-GFP* strain were inoculated onto the rice sheaths. At 24 hpi the fluid in the conidial suspension was replaced with either sterile H_2_O (Control), 8 mg/mL sucrose, 50 mg/mL glucose, or 1.5 mg/mL G6P (Glucose-6-phosphate). Confocal microscopy was carried out at 30, 48, and 72 hpi. The right panels show enlarged view of the boxed region in the left panel. Scale bar equals 5 μm.

**Figure 7.**
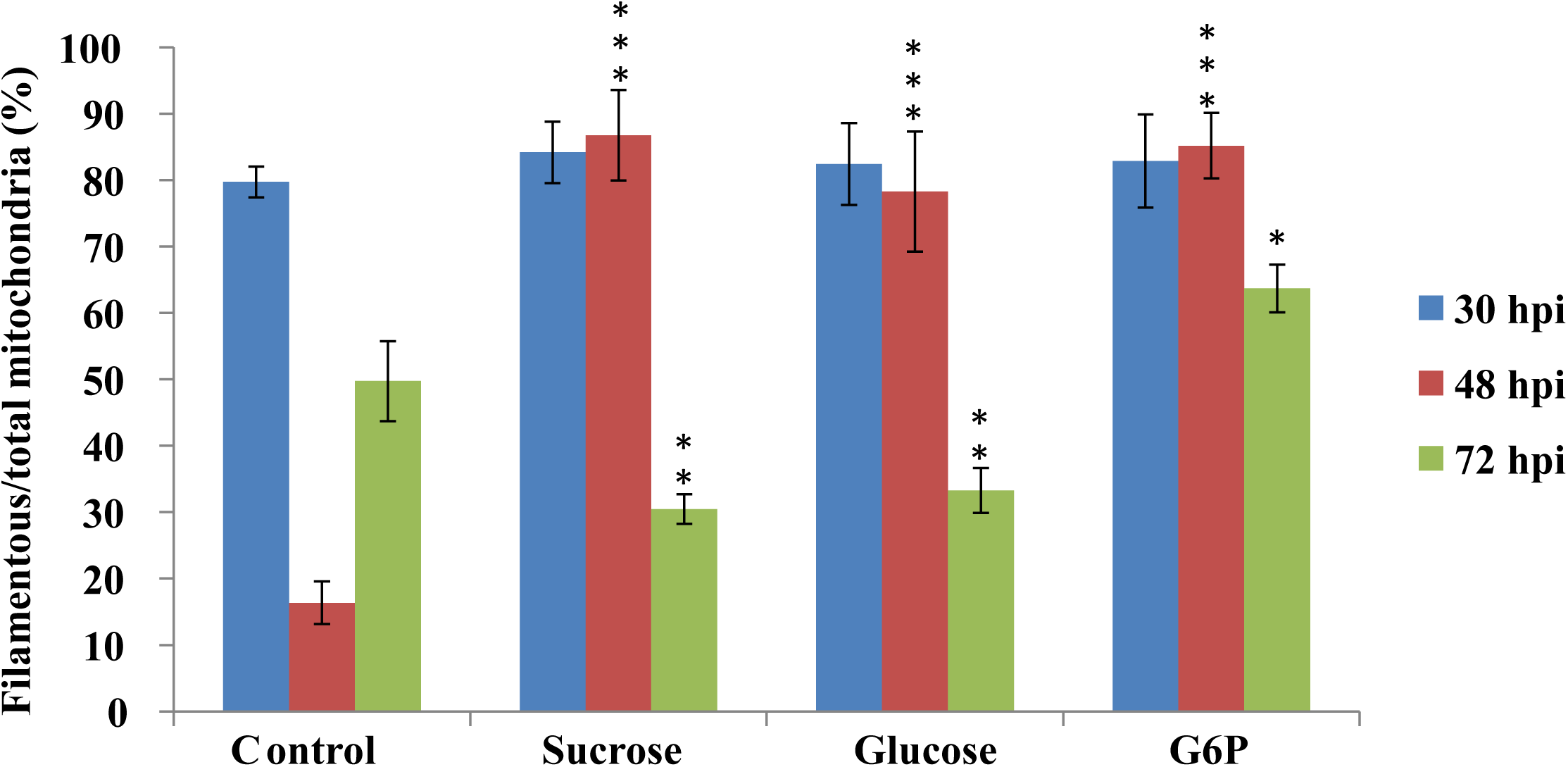
Mitochondrial morphology with or without exogenous sucrose, glucose, or G6P. Values represent the mean ± SD from three independent experiments. ***, P<0.001; **, P<0.005; *, P<0.01 in comparison to control (H_2_O) at the same time points. Sample size is more than 200 appressoria penetration sites per analysis.

### Mitophagy is necessary for blast infection

As shown in Figure 1, 3, 6, and 8d, the vacuolar localization of Mito-GFP (mitochondrial marker) was observed at 48 hpi and 72 hpi (Figure S6), indicating that mitophagy is likely induced to degrade mitochondria during the initial stages of establishment of the blast disease. Our previous study showed that MoAtg24 is specifically required for mitophagy and is necessary for proper asexual differentiation (He et al., 2013). To determine whether mitophagy plays any role during infection, the pathogenicity of *Moatg24*Δ was tested using rice seedling infection assays. Compared to the WT, which caused the characteristic spindle-shaped blast lesions with grey centres, the *Moatg24*Δ showed highly reduced pathogenicity in rice (Figure 8a). Typical blast disease lesions were not elaborated in the susceptible rice cultivar inoculated with *Moatg24*Δ conidia, while only small lesions were occasionally evident (Figure 8a).

**Figure 8.**
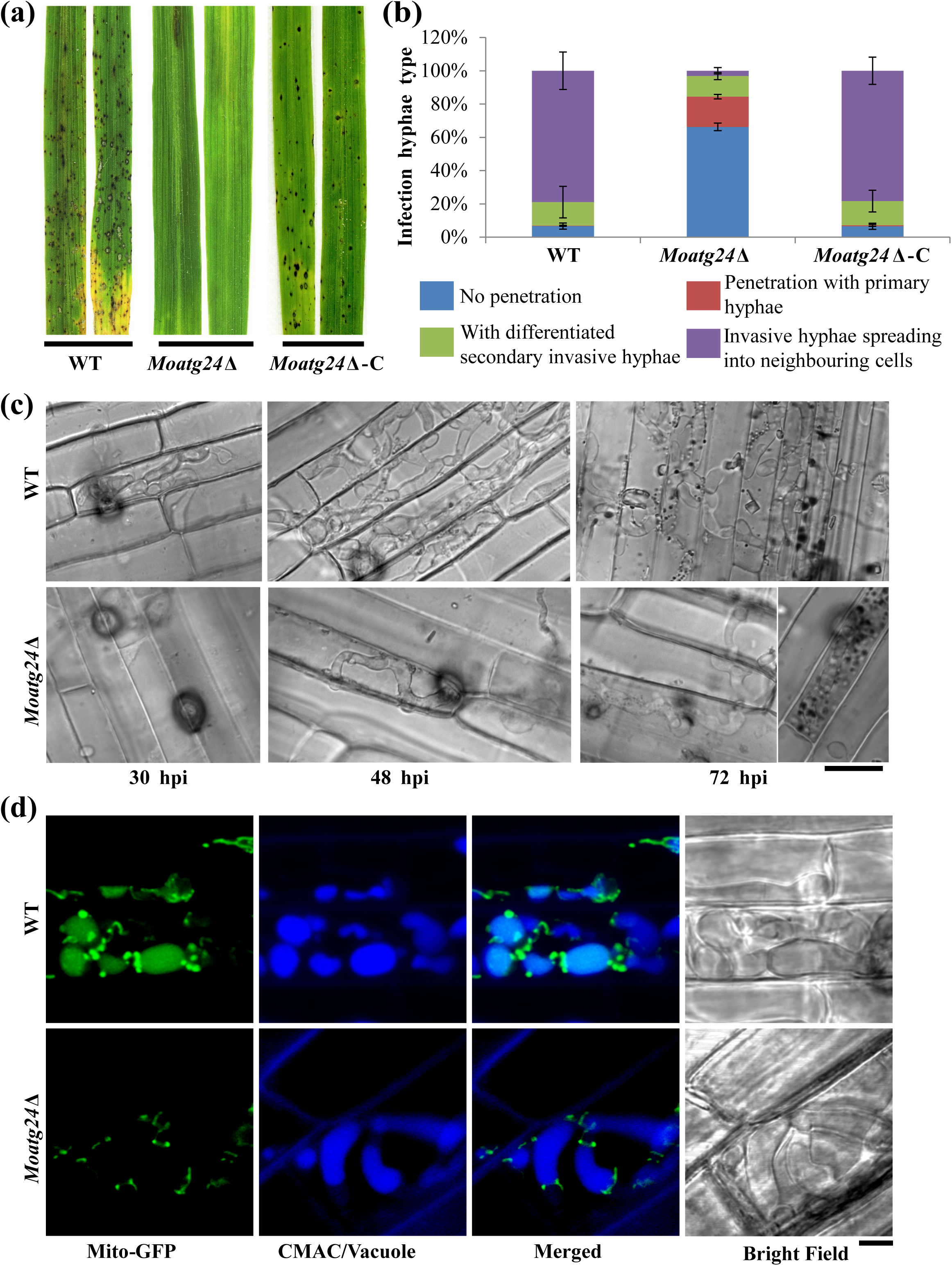
MoAtg24-mediated mitophagy is necessary for *M. oryzae* infection. (a) Loss of *MoATG24* gene leads to reduction in pathogenicity. Blast infection assays of wild type (WT), *Moatg24*Δ, or *Moatg24*Δ complementation strain (*Moatg24*Δ-C) were performed using rice seedlings (*Oryza sativa* L., cultivar CO39). Images were taken at 7 dpi. (b) Developmental defects in the invasive hyphae of *Moatg24*Δ strain. The invasive hyphae in rice sheath cells were quantified as descripted in Figure 2c. Data represents the mean ± SD from three independent experiments. More than 200 appressoria from each indicated strain were assessed each time. (c) Invasive hyphal growth in WT and *Moatg24*Δ strains at 30, 48, and 72 hpi. Scale bar = 12 μm. (d) MoAtg24 is required for mitophagy during *M. oryzae* infection. WT or *Moatg24*Δ strain expressing *Mito-GFP* was inoculated into rice sheaths for 60 h. The vacuoles in invasive hyphae were visualized by staining with CMAC. Scale bar = 2.5 μm.

To understand the differences between infection by *Moatg24*Δ and WT conidia, invasive hyphae were observed under the microscope at 30 hpi, 48 hpi, and 72 hpi. At 40 hpi, nearly 90% of WT appressoria successfully penetrated the rice sheath. By contrast, less than 20% of *Moatg24*Δ appressoria were capable of invading the rice sheath (Figure 8b; P<0.001). At 48 hpi, although the penetration rates of appressoria in WT and *Moatg24*Δ were comparable, the secondary invasive hyphae were highly reduced in the *Moatg24*Δ (less than 2%) compared to WT (around 80%) (Figure 8b; P<0.001). At 72 hpi, the difference in invasive hyphae in *Moatg2*4Δ and WT was more pronounced. The invasive hyphae of WT had successfully spread into 5 to 7 rice cells, whereas the invasive hyphae of *Moatg24*Δ were mainly restricted to the first invaded cells in the rice epidermis (Figure 8c). In rare cases, the invasive hyphae of *Moatg24*Δ could be found within the neighbouring cells surrounding the primary infected rice epidermal cell. Based on these results, we inferred that the highly reduced pathogenicity of *Moatg24*Δ is a result of lack of spread of invasive hyphae from the site of host entry/invasion.

Since MoAtg24 is essential for mitophagy, and the *Moatg24*Δ showed highly reduced invasive growth in this study, it became important to assess and confirm whether MoAtg24-based mitophagy occurs naturally during blast infection. As shown in Figure 8d, Mito-GFP signal could be detected in vacuoles (CMAC staining), at 60 hpi in invasive hyphae (Figure 8d, upper panel). In contrast, such Mito-GFP signal did not colocalize with the vacuoles in invasive hyphae in *Moatg24*Δ (Figure 8d, lower panel), indicating that mitophagy during infection-related growth of *M. oryzae* is blocked in *Moatg24*Δ mutant. Considering that the *Moatg24*Δ mutant is defective in invasive growth and failed to form blast lesions, these results showed that *MoATG24*-mediated mitophagy plays a critical role in the infection process of *M. oryzae.*

Taken together, our data support that mitochondrial dynamics and mitophagy are important intermediate events between nutrient sensing and homeostasis in *M. oryzae* leading to the establishment and extent of the devastating blast disease in rice.

## DISCUSSION

The adaptation of the mitochondrial fusion and fission to cellular demands is critical for a number of important physiological processes (Zemirli & Morel, 2018). The defects in mitochondrial dynamics cause severe physiological consequences and lead to a variety of dysfunctions (Delettre et al., 2000, Zemirli & Morel, 2018, Kijima et al., 2005), thus highlighting that fusion and fission process must be tightly controlled. In this study, the mitochondrial dynamics were first observed during the early stages of the infection cycle of *M. oryzae*. Interestingly, we uncovered a unique filamentous-punctate-filamentous transition cycle in mitochondrial morphology during *in planta* growth. We demonstrated that the key regulators of mitochondrial fusion and fission are essential for proper mitochondrial dynamics and invasive growth in *M. oryzae*. These results suggest that mitochondrial fusion and fission are tightly controlled during blast infection; and that such sequential change in organellar morphology is important for pathogenesis of *M. oryzae*.

Mitochondrial fragmentation could be triggered by multiple environmental factors or stressors. The blast pathogen generally encounters host defense, oxidative stress, and metabolic stress that may trigger such mitochondrial fragmentation. We propose that carbon source depletion is one of the important factors triggering mitochondrial fragmentation during *in planta* growth of *M. oryzae*. Firstly, mitochondrial fragmentation was observed in the invasive hyphae in heat-killed rice sheath, which is incapable of mounting the defense response. Secondly, exogenous antioxidants did not inhibit or delay such fragmentation processes during infection. Carbon sources (such as glucose or sucrose) or the important metabolic intermediate (G6P) delayed mitochondrial fragmentation, whereas prolonged nutrient starvation induced the breakdown of mitochondrial network in *M. oryzae* (Figure S7). In addition, we found that NH_4_NO_3_ treatment did not change the mitochondrial morphology (Figure S8), indicating that nitrogen starvation is likely not an important factor that leads to mitochondrial fragmentation during blast infection. Strigolactone is a plant hormone which is associated with mitochondrial biogenesis, fission, fusion, spore germination and hyphal branching in some fungal genera (Besserer *et al.*, 2006). Strigolactone (GR24) treatment did not inhibit the mitochondrial fragmentation processes during *Magnaporthe* infection (Fig S8).

*M. oryzae* initially acquires nutrients from living host cells, but switches to the necrotrophic killing phase to acquire nutrients from dead tissues between 48 and 72 hpi (Figure S9c). During the transition to necrotrophy, the filamentous invasive hyphae of *M. oryzae* maintain viability as the fungal lifestyle changes and lesion development when host cell death is occurring (Kankanala *et al.*, 2007, Fernandez & Orth, 2018, Jones *et al.*, 2016). *M. oryzae* thus needs to adapt to and overcome nutrient stress prior to switching to the necrotrophic phase. Our results showed that glucose or sucrose supplementation promotes proliferation of invasive hyphae and decreases the cell death in rice during early infection of *M. oryzae* (Figure S9). These results suggest that carbon source depletion occurs during infection and is likely a major factor which triggers the biotrophy-necrotrophy transition. However, carbon starvation and then carbon source acquisition from dead plant tissues may not be the only factor that triggers mitochondria fragmentation and rebuilding of the network, since addition of glucose every 6 h after 24 hpi, simply delayed the fragmentation of mitochondria (Figure 6, 7). It is possible that other signals cooperate with carbon homeostasis machinery to regulate and control the mitochondrial morphology/function during the blast infection in rice. In conclusion, our study suggests that carbon source depletion with other factor(s) trigger(s) mitochondrial fragmentation and biotrophic-necrotrophic phase switch during infection of *M. oryzae*.

In this study, we found that mitophagy is induced along with precise mitochondrial fragmentation during rice blast. Furthermore, mitophagy plays a critical role in invasive growth of *M. oryzae* in response to energy demands and nutrient homeostasis. It has been suggested that mitophagy requires efficient fission to separate out damaged or unwanted mitochondria to fit into the autophagosomes (Mao & Klionsky, 2013). Thus, it is possible that mitochondrial fragmentation together with ensuing mitophagy is the strategy employed by *M. oryzae* to separate and degrade the damaged/excess mitochondria in order to protect itself in the hostile environment *in planta*.

In conclusion, our study revealed that a unique filamentous-punctate-filamentous cycle in mitochondrial morphology controlled by fission and fusion machinery is important for pathogenesis of *M. oryzae*. Such morphological transitions likely couple with nutrient homeostasis (particularly carbon source) and biotrophy-to-necrotrophy switch during *M. oryzae* infection. In addition, mitophagy regulates the precise turnover of mitochondria, and plays a critical role during the initiation of the devastating blast disease in rice.

## EXPERIMENTAL PROCEDURES

### Fungal strains and culture media

The *M. oryzae* WT strain B157 (field isolate, *mat1-2*) was a kind gift from the Indian Institute of Rice Research (Hyderabad, India). The *M. oryzae* strains *Mito-GFP, Moatg24*Δ, and *Moatg24*Δ-C have been described in our previous reports (Patkar et al., 2012, Ramos-Pamplona & Naqvi, 2006, He et al., 2013).

*M. oryzae* strains were grown on prune agar (Yeast extract 1 g/L, lactose 2.5 g/L, sucrose 2.5 g/L, prune juice 40 mL/L, agar 20 g/L, pH 6.5) medium at 28°C in the dark for 2 days, followed by growth under continuous light for 5 days to collect conidia for infection assay. Mutants generated by *Agrobacterium tumefaciens-*mediated transformation (ATMT) were selected on either Complete medium (CM: Casein Hydrolysate 6 g/L, Sucrose 10g/L, Yeast Extract 6 g/L, Agar 20 g/L) containing Hygromycin (250 μg/ml) or Basal medium (BM: Asparagine 2.0 g/L, Yeast Nitrogen Base 1.6 g/L, NH_4_NO_3_ 1.0 g/L, Glucose 10 g/L, Agar 20g/L, pH 6.0) with Chlorimuron-ethyl (50 μg/ml) or with Ammonium glufosinate (50 μg/ml).

### Construction of *Modnm1*Δ and *Mofzo1*Δ strains and complementation analyses

The *MoDNM1* gene (*MGG_06361*) deletion mutant was generated using the standard one-step gene replacement strategy. Briefly, about 1 kb (kilobase) of 5’ UTR and 3’ UTR regions were PCR amplified and ligated sequentially to flank the *ILV2*^*SUR*^ sulfonylurea-resistance cassette in pFGL820 (Addgene, 58221) (Figure S2a). The following primers were used to amplify the 5’ and 3’ UTR of the *MoDNM1* gene: Dnm1-5F (5’-GAGAGTGTT <u>GAATTC</u> CTCACGGGATGGGCTTCTG-3’) Dnm1-5R (5’-GAGAGTGTT <u>GGTACC</u> GGCGAAAATCGGTTCCGTGGTC-3’), Dnm13-F (5’-GAGAGTGTT <u>GTCGAC</u> TGAAGCTGTTTGCGCCATG-3’), and Dnm13-R (5’-GAGAGTGTT <u>GCATGC</u> TACCTATGATCAGCCCGC-3’). Underlined sequences are restriction sites introduced for cloning purpose. The final plasmid construct was confirmed by sequencing and subsequently introduced into the *Mito-GFP* strain by ATMT to replace the *MoDNM1* gene (Yang & Naqvi, 2014). All the correct transformants in this study were ascertained by locus-specific PCR and/or Southern blot analysis (Figure S2b, S3b). For complementation analysis, the full length genomic copy with promoter of *MoDNM1* was amplified with MoDnm1-F (5’-AATT <u>GAATTC</u> GTTGAGCAGGCCGAGCGAC-3’) and MoDnm1-R (5’-AATT <u>GAATTC</u> CACTGGCATTTGATTACGCAAGG-3’) inserted into pFGL822 (Addgene, 558226) and introduced into the *Modnm1*Δ strain.

For generating the plasmid vector for *MoFZO1* (*MGG_05209*) deletion, about 1 kb of 5’ UTR and 3’ UTR regions were PCR amplified and ligated sequentially to flank the *phosphinothricin acetyl transferase* gene cassette in pFGL822 (Figure S3). The following primers were used to amplify the 5’ and 3’ UTR of the Mo*Fzo1* gene: Fzo1-5F (5’-GAGAGTGTT <u>GAATTC</u> ACTCGGCCGCGATACGCTGC-3’), Fzo1-5r (5’-GAGAGTGTT <u>GGATCC</u> GTGATCGATTTCGTCCAGTC-3’), Fzo1-3F (5’-GAGAGTGTT <u>CTGCAG</u> GCAGAACCATCCTCGTCGTC-3’), and Fzo1-3r (5’-GAGAGTGTT <u>AAGCTT</u> CCTGGCGGCGGCGACATCAAC-3’). The final plasmid was introduced into the *Mito-GFP* strain by ATMT to replace the *MoFZO1* gene. The complementation fragment, which contains the full length genomic copy with promoter of *MoFZO1* gene, was amplified with MoFzo1-F (5’-AATT <u>GGATCC</u> GGCTGTCTGCGTGATCCCTG-3’) and MoFzo1-R (5’-AATT <u>TCTAGA</u> GCTGTGGAGCGAGGAGCAGG-3’) and inserted into pFGL899 to complement the *Mofzo1*Δ strain (Yang & Naqvi, 2014).

### Infection assays

For blast infection assay, conidial suspension (10^6^/mL) with 0.01% gelatine was sprayed on 21 day old rice seedlings (*Oryza sativa* L., cultivar CO39) and incubated in a growth chamber (16 h light/d, 22°C and 90% humidity). Blast disease in infection assays were assessed and recorded by scanning the leaves at 7 days post inoculation. The blast infection assays were repeated at least three times.

For the host penetration and *in planta* invasive hyphal development assay, healthy rice seedlings (CO39) at the age of 4 weeks were selected for sheath preparation. Conidial suspension (5×10^4^/ mL) were inoculated onto rice sheath and incubated on the sterile wet tissue paper in the 90 mm Petri dish. Then the petri dishes with inoculated rice sheaths were transferred into the growth chamber with a photoperiod of 16 h: 8 h light: dark cycle at 25°C. The inoculated sheath was trimmed manually and observed by using an Olympus BX51 wide field microscope or with a laser scanning confocal microscope at selected time points.

To prepare heat-killed rice sheaths, the fresh rice sheaths were immersed into sterile water at 70°C for 25 min (Shipman *et al.*, 2017). The heat-killed rice sheath has the physical structures of cells, while the abilities of host response to fungal infection are lost.

### Carbon sources, antioxidant, and Mdivi-1 treatments

For treatments with excess carbon sources, the conidia from the tested strains were inoculated on to rice sheath and incubated in growth chamber. At 24 hpi (hours post inoculation), the water on the rice sheath was removed, and then following solutions were applied to the sheath: 8 mg/mL sucrose, 50 mg/mL glucose, 1.5 mg/mL G6P (Glucose-6-phosphate, Sigma-Aldrich), 2.5 mM GSH (L-Glutathione reduced, Sigma-Aldrich), 40 mM NAC (N-acetyl cysteine, Sigma-Aldrich), or 10 mM Mdivi-1 (Selleck). The rice sheaths were incubated in the growth chamber until observation. These experiments were repeated thrice.

### Vacuolar staining

The infected rice sheaths were incubated with CellTracker™ Blue CMAC Dye (7-amino-4-chloromethylcoumarin, Molecular Probes, C2110) at a final working concentration of 10 μM for 2 h at 37 °C. The sample was washed with water prior to microscopic observation.

### Live cell imaging and image processing

Live cell epifluorescence microscopy was performed with a Zeiss LSM 700 inverted confocal microscope (Carl Zeiss, Inc) using a Plan-Apochromat 63 (NA=1.40) Oil immersion lens. EGFP (Enhanced GFP) and CMAC excitation were performed at 488 nm (Em. 505-530 nm) and 405 nm (Em. 430-470 nm) respectively. For *in planta* invasive hyphal development observation, z-stack that consisted of 0.5 μm-space sections was captured for each appressorium penetration site. Image processing was processed in Image J program which was downloaded from National Institutes of Health (http://rsb.info.nih.gov/). The maximum projection of z-stack was obtained by Z projection with max intensity in Image J. 3-D reconstruction, visualization, and analysis were performed in Bitplan Imaris with filament and spots program (Zurich, Switzerland). For figure preparation, the images were arranged in Adobe Illustrator CS6.

## CONFLICT OF INTEREST

There is no conflict of interest.

### ACKNOWLEDGEMENTS

We thank the Fungal Pathobiology group at TLL for useful discussion and suggestions. We thank Gu Keyu and Amit Anand for image analyses. This study was supported by the Chinese Academy of Agricultural Sciences under “Elite Youth” program and Agricultural Sciences and Technologies Innovation Program, and the Zhejiang Provincial Natural Science Foundation of China (LQ19C140004). National Research Foundation, Singapore (Prime Minister’s Office, NRF-CRP7-2010-02), and intramural grants from the Temasek Life Sciences Laboratory, Singapore.

Naweed I. Naqvi, Yanjun Kou, and Yunlong He planned and designed the research. Yanjun Kou, Yunlong He, Yazhou Shu, Jiehua Qiu, Fan Yang, and Yizhen Deng performed experiments, conducted fieldwork, analysed data etc. Yanjun Kou, Yunlong He, and Naweed I. Naqvi wrote the manuscript. Yanjun Kou and He Yunlong contributed equally.

## SUPPORTING INFORMATION LEGENDS

**Figure S1** Mitochondrial morphology during appressorium formation in *M. oryzae*.

**Figure S2** Generation and verification of *Modnm1*Δ mutant.

**Figure S3** Generation and verification of *Mofzo1*Δ mutant.

**Figure S4** Oxidant treatment induces mitochondrial fragmentation.

**Figure S5** Antioxidant treatment does not delay mitochondrial dynamics during invasive growth.

**Figure S6** The vacuolar localization of Mito-GFP (mitochondrial marker) during invasive growth.

**Figure S7** Prolonged nutrient starvation induces mitochondrial fragmentation and mitophagy.

**Figure S8** NH_4_NO_3_ or GR24 treatment did not change the mitochondrial fragmentation during infection by *M. oryzae*.

**Figure S9** Addition of glucose or sucrose promotes spread of invasive hyphae and decreases the cell death in rice during early infection by *M. oryzae*.

