## Supplementary material for "Mitochondrial dynamics and mitophagy are necessary for proper invasive growth in Rice Blast": Figure S

### Fig. S1

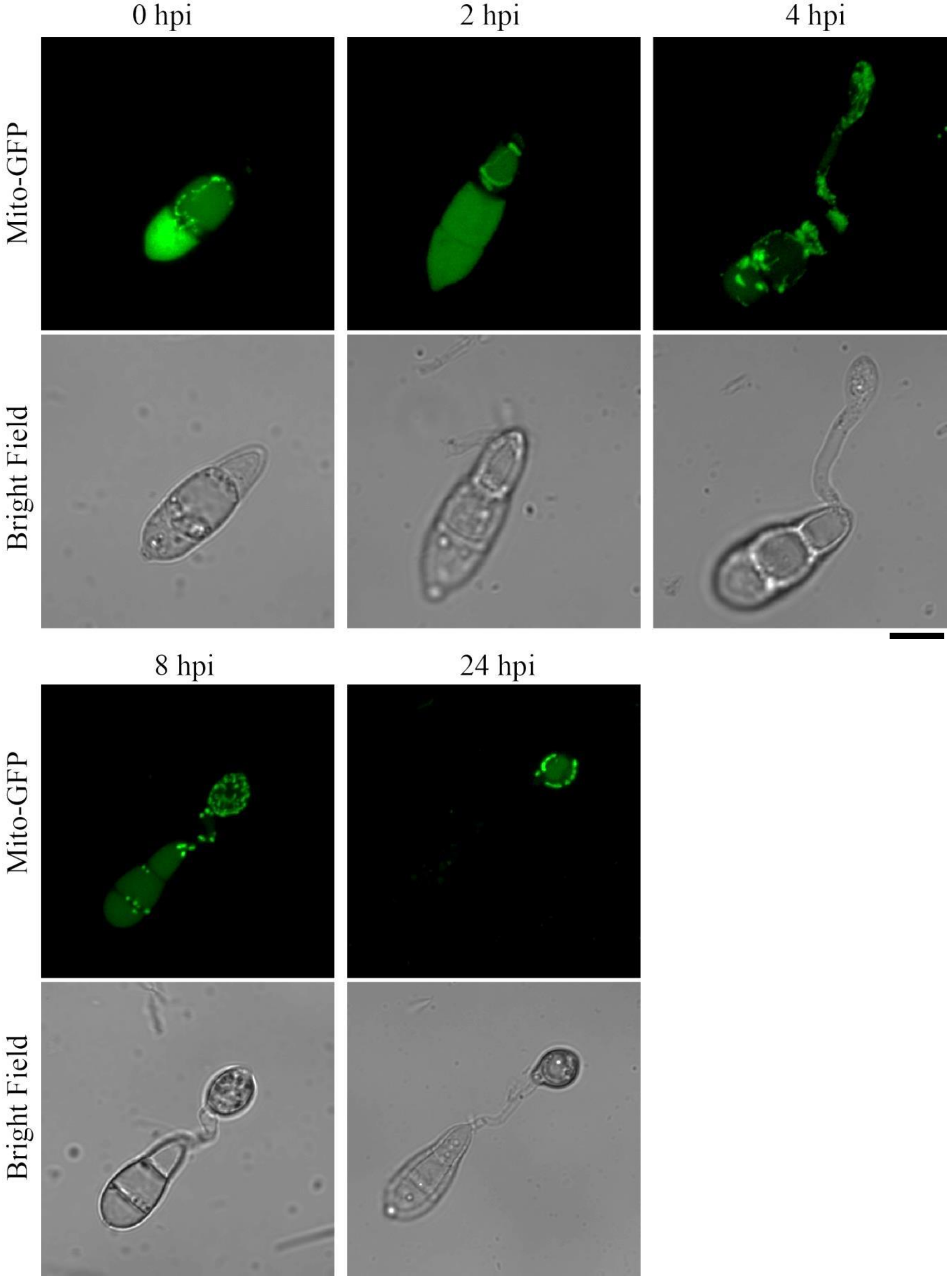


**Fig. S1** Mitochondrial morphology during appressorium formation in *M. oryzae*. The conidial suspension of the *Mito-GFP* strain was inoculated on the inductive surface. Confocal microscopy was carried out at 0, 2, 4, 8, and 24 hpi. Scale bar: 10 μm

**Fig. S2**

(a)

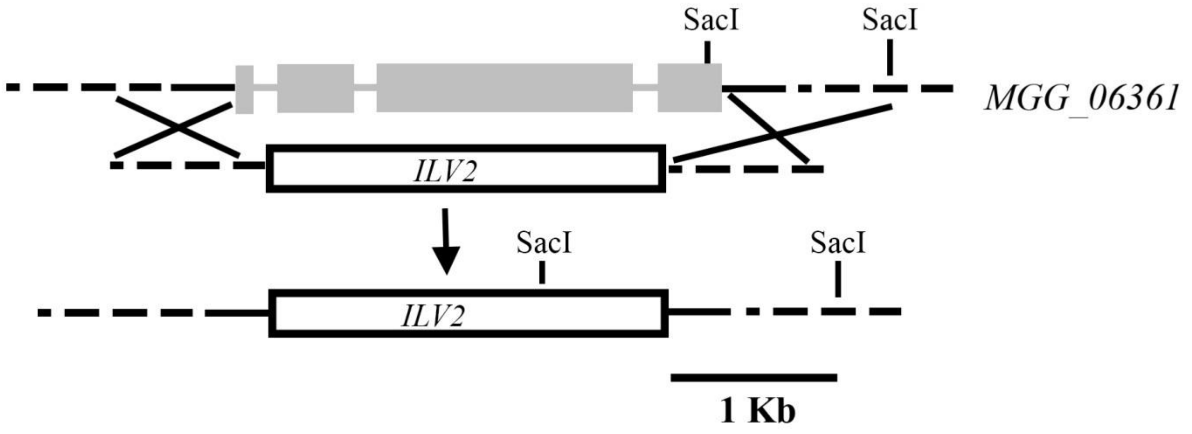


(b)

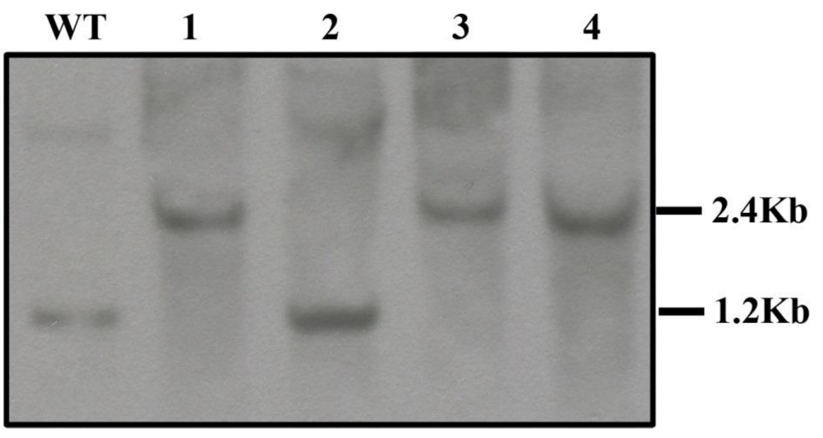


**Fig. S2** Generation and verification of *Modnm1*Δ mutant. (a) Schematic representation of the *MoDNM1* (*MGG_06361*) locus with flanks (dashed lines) in *M. oryzae*. Exons (grey solid bars) and introns (grey lines) are drawn to scale. The *MoDNM1* coding region was replaced with *ILV2*, which represents sulfonylurea-resistant *ILV2*$^{SUR}$ variant gene cassette, by homologous recombination. (b) Southern blot analysis was conducted to confirm the *Modnm1*Δ mutant. *Sac*I-digested genomic DNA from the WT or *Modnm1*Δ mutant was probed with denoted 1kb flank in (a). The appearance of the 2.4 kb band in the *Modnm1*Δ mutant, with the concomitant loss of the WT 1.2 kb in *MoDnm1* locus, is indicative of the correct gene replacement event.

### Fig. S3

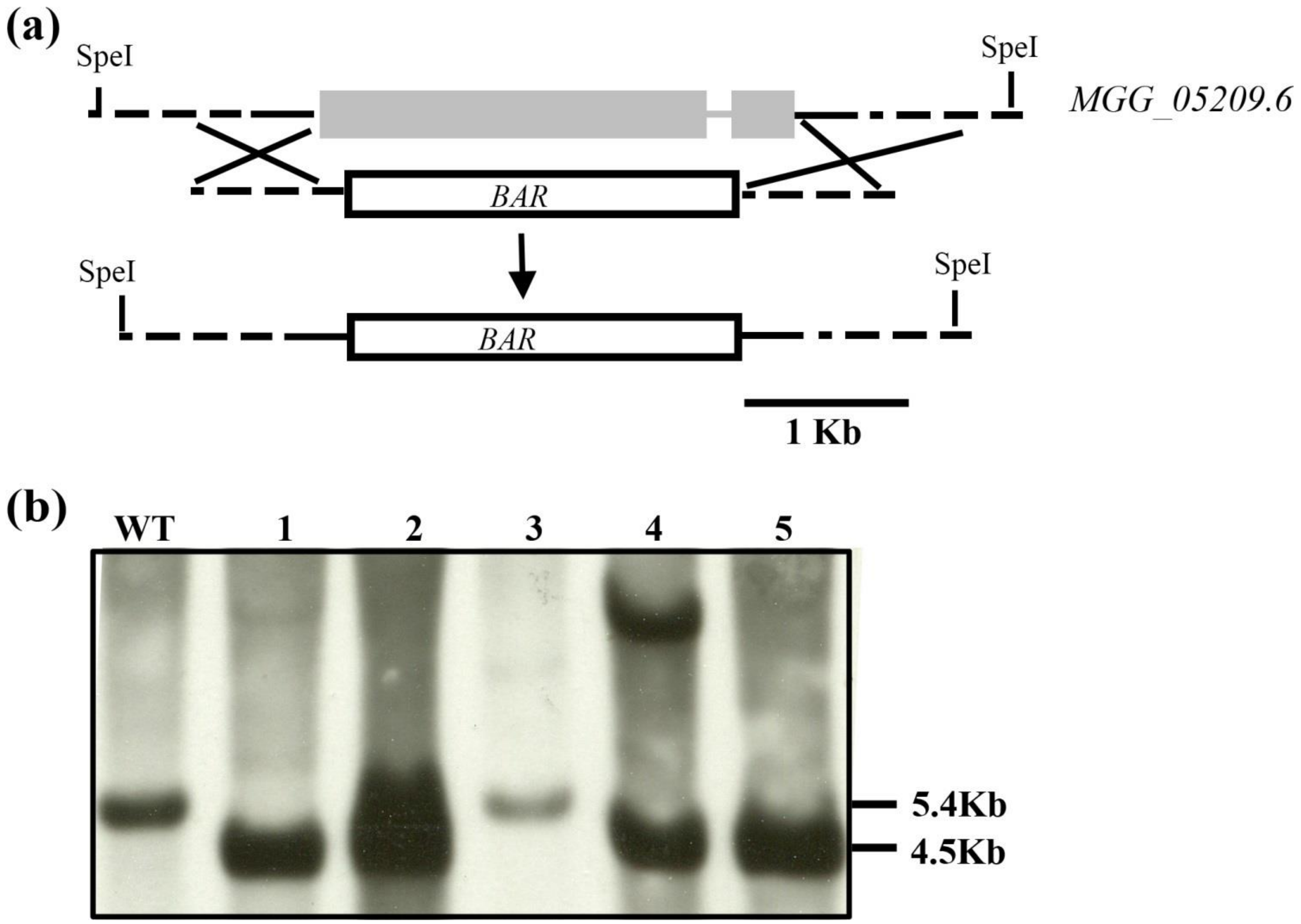


**Fig. S3** Generation and verification of *Mofzo1*Δ mutant. (a) Schematic representation of the *MoFZO1* (*MGG_05209*) locus with flanks (dashed lines) in *M. oryzae*. Exons (grey solid bars) and introns (grey lines) are drawn to scale. The *MoFZO1* coding region was replaced with *BAR*, which represents *Phosphinothricin acetyl transferase* gene cassette, by homologous recombination. Scale bar denotes 1 kb. (b) Southern blot analysis was used to confirm the *Mofzo1*Δ mutant. *Spe*I-digested genomic DNA from the WT and *Mofzo1*Δ mutant were probed with the denoted 1kb flank in (a). The WT 5.4 kb *MoFZO1* locus was lost while a 4.5 kb band was detected in *Mofzo1*Δ mutant, which was diagnostic of the correct gene replacement event.

### Fig. S4

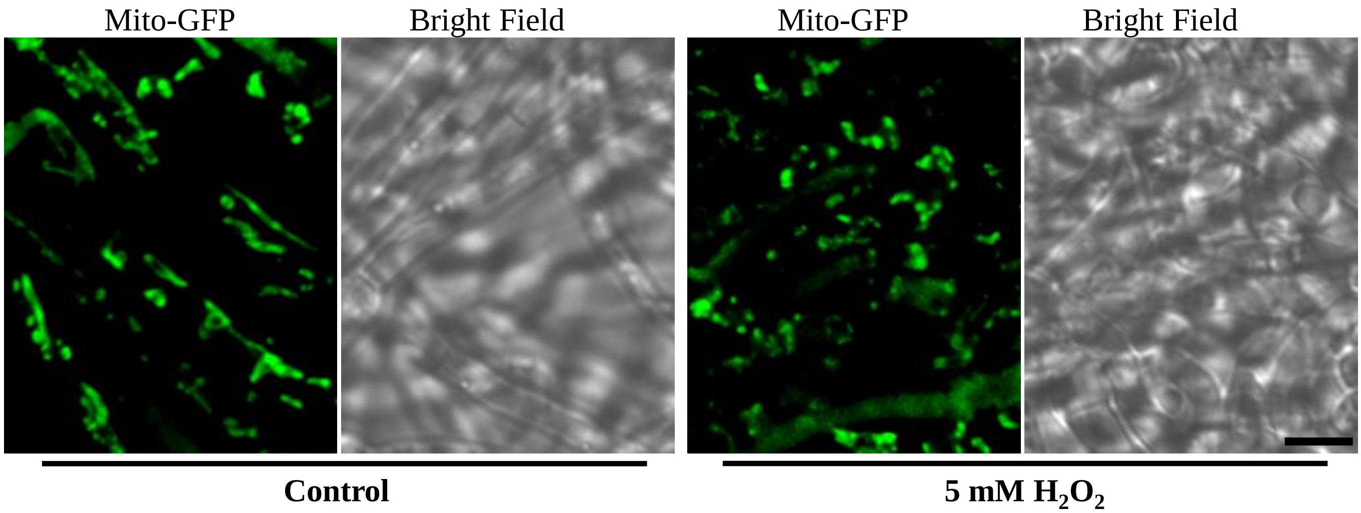


**Fig. S4** Oxidant treatment induces mitochondrial fragmentation. The *Mito-GFP* strain was grown in liquid CM for 2 d followed by inoculation in for CM with 5 mM $H_2O_2$ for 12 h. Scale bar = 5 μm.

Fig. S5

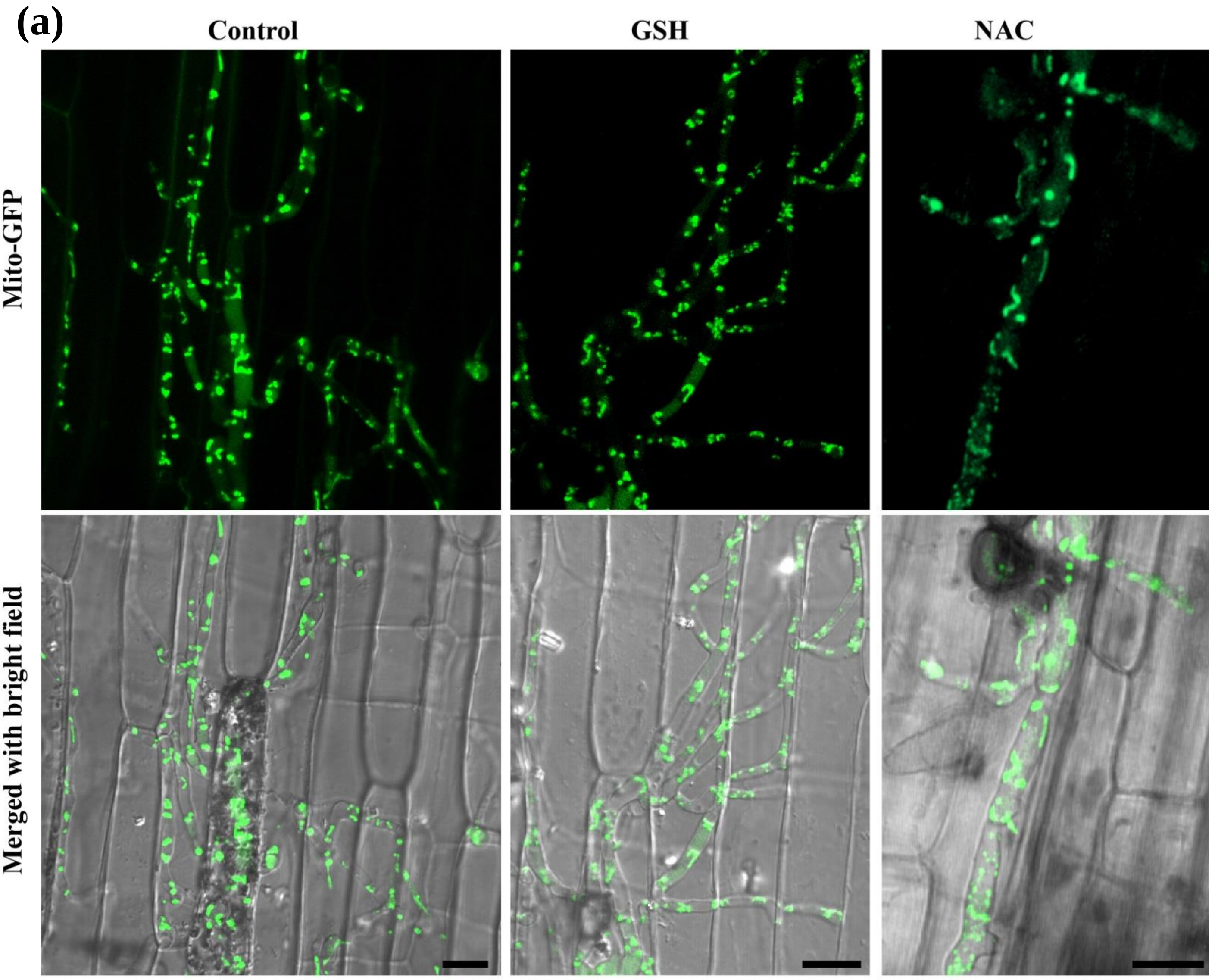

(a)
Control
GSH
NAC
Mito-GFP
Merged with bright field


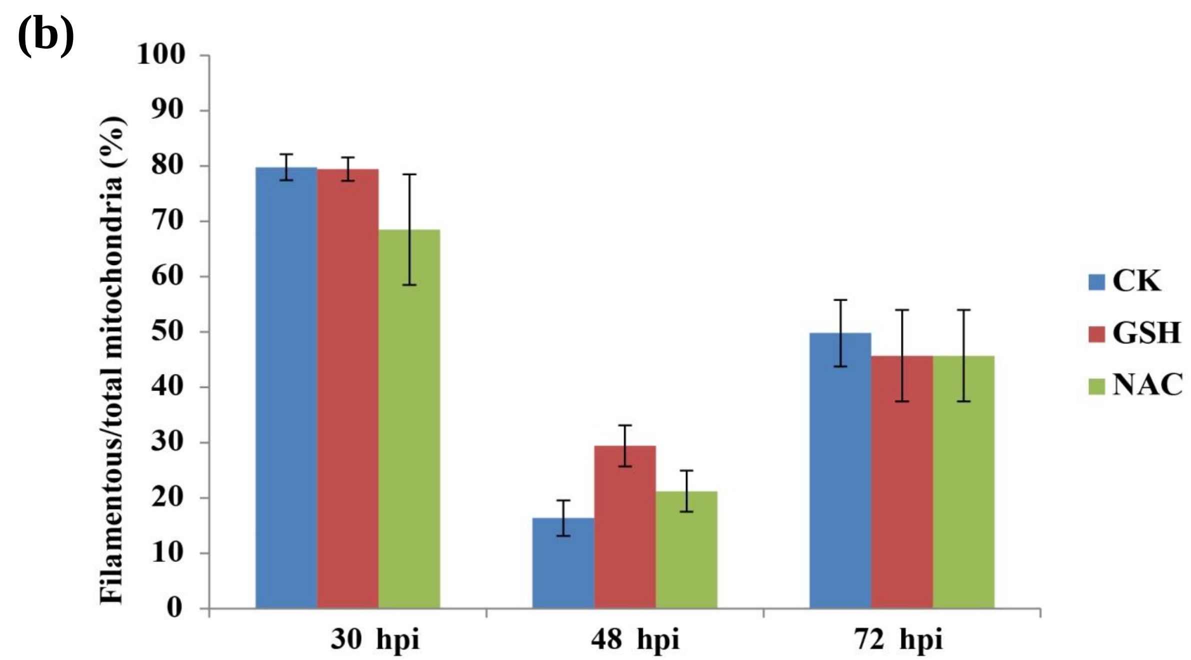

(b)
Filamentous/total mitochondria (%)
100
90
80
70
60
50
40
30
20
10
0
30 hpi
48 hpi
72 hpi
CK
GSH
NAC

**Fig. S5** Antioxidant treatment does not delay mitochondrial dynamics during invasive growth. (a) Confocal microscopy image of the *Mito-GFP* strain in rice sheath cells in present of GSH or NAC. A conidial suspension of the *Mito-GFP* strain was inoculated on the rice sheath for 24 h before 2.5mM GSH or 40 mM NAC was added. Confocal microscopy was carried out at 48 hpi. Scale bar: 10 μm. (b) Quantitative analysis of mitochondria of the *Mito-GFP* strain in rice sheath cells with GSH or NAC treatment. Values represent the mean ± SD from three independent experiments. Sample size is more than 50 appressoria penetration sites per analysis.

### Fig. S6

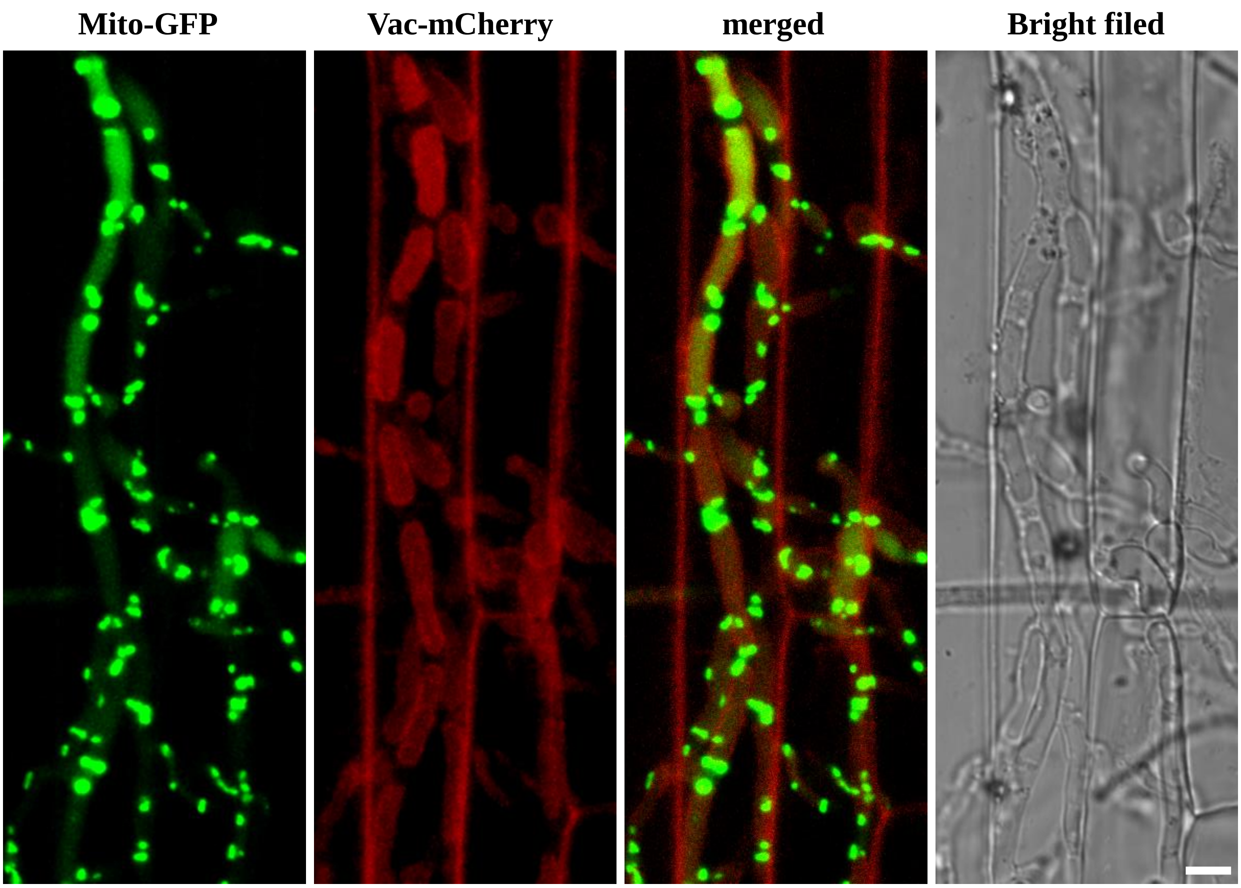


**Fig. S6** The vacuolar localization of Mito-GFP (mitochondrial marker) during invasive growth. Scale bar: 5 µm. The *V-type proton ATPase subunit A* (*MGG_03947*) gene was tagged with *mCherry* to highlight the vacuole of *M. oryzae* at 48 hpi.

### Fig. S7

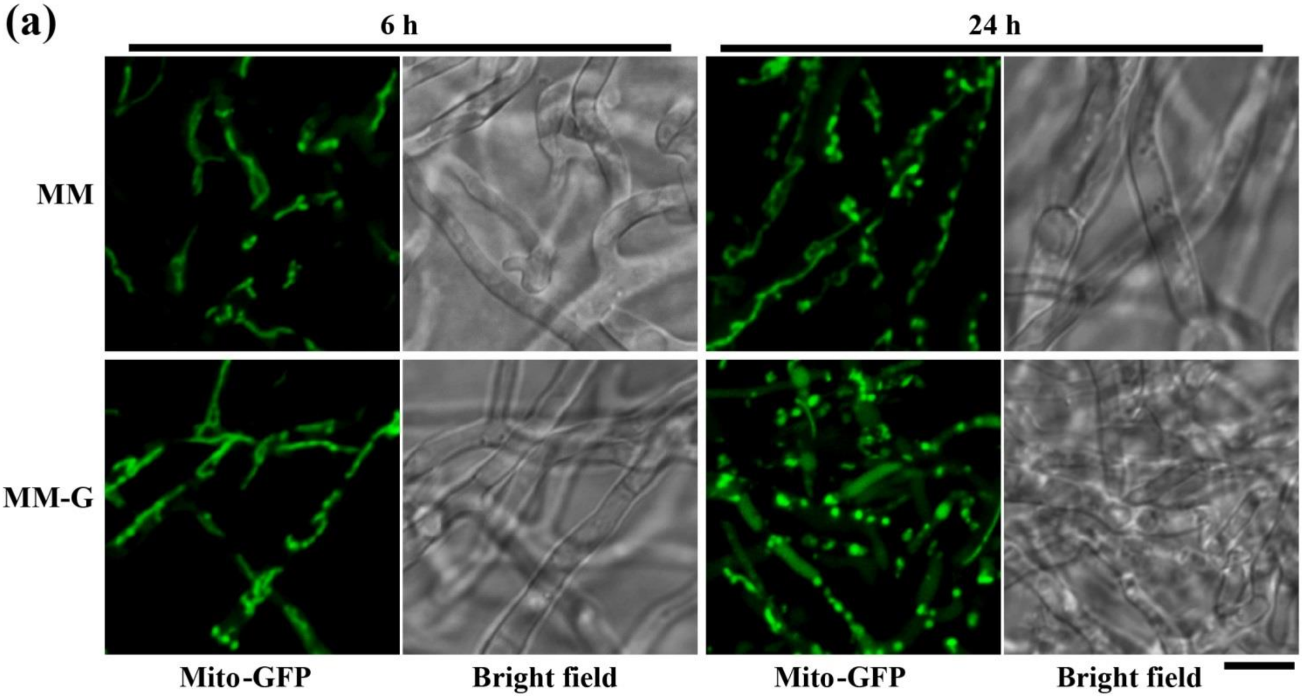
(a)
6 h
24 h
MM
MM-G
Mito-GFP
Bright field
Mito-GFP
Bright field

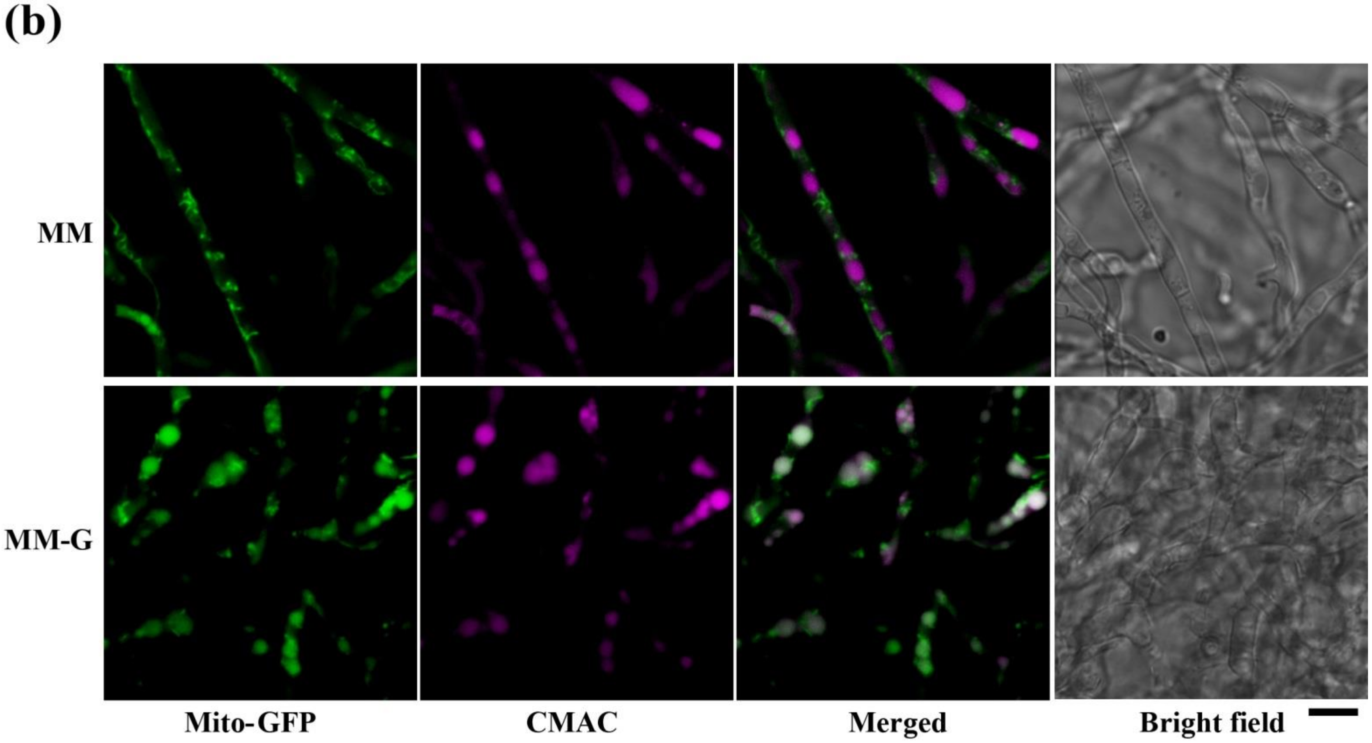
(b)
MM
MM-G
Mito-GFP
CMAC
Merged
Bright field

**Fig. S7** Prolonged nutrient starvation induces mitochondrial fragmentation and mitophagy. (a) Mitochondrial morphology under depletion of carbon source starvation. The *Mito-GFP* strain was grown in liquid CM for 2 days followed by inoculation in liquid MM (Minimal medium; 6 g/L $NaNO_3$, 0.5 g/L $MgSO_4$, 0.5 g/L KCl, 1.5 g/L $KH_2PO_4$, 10 g/L glucose, 0.1% (v/v) trace elements, pH 6.5) or MM-G (Minimal medium lacking glucose; 6 g/L $NaNO_3$, 0.5 g/L $MgSO_4$, 0.5 g/L KCl, 1.5 g/L $KH_2PO_4$, 0.1% (v/v) trace elements, pH 6.5) for 6 h or 24 h. The fragmented mitochondria are trafficked to the vacuoles for degradation under the carbon source starvation for 24 h. (b) Mitophagy is induced by nutrient starvation for 24 h. The vacuoles in invasive hyphae were stained by CMAC and rendered in pseudo-color using Image J. Scale bar = 5 μm.

### Fig. S8

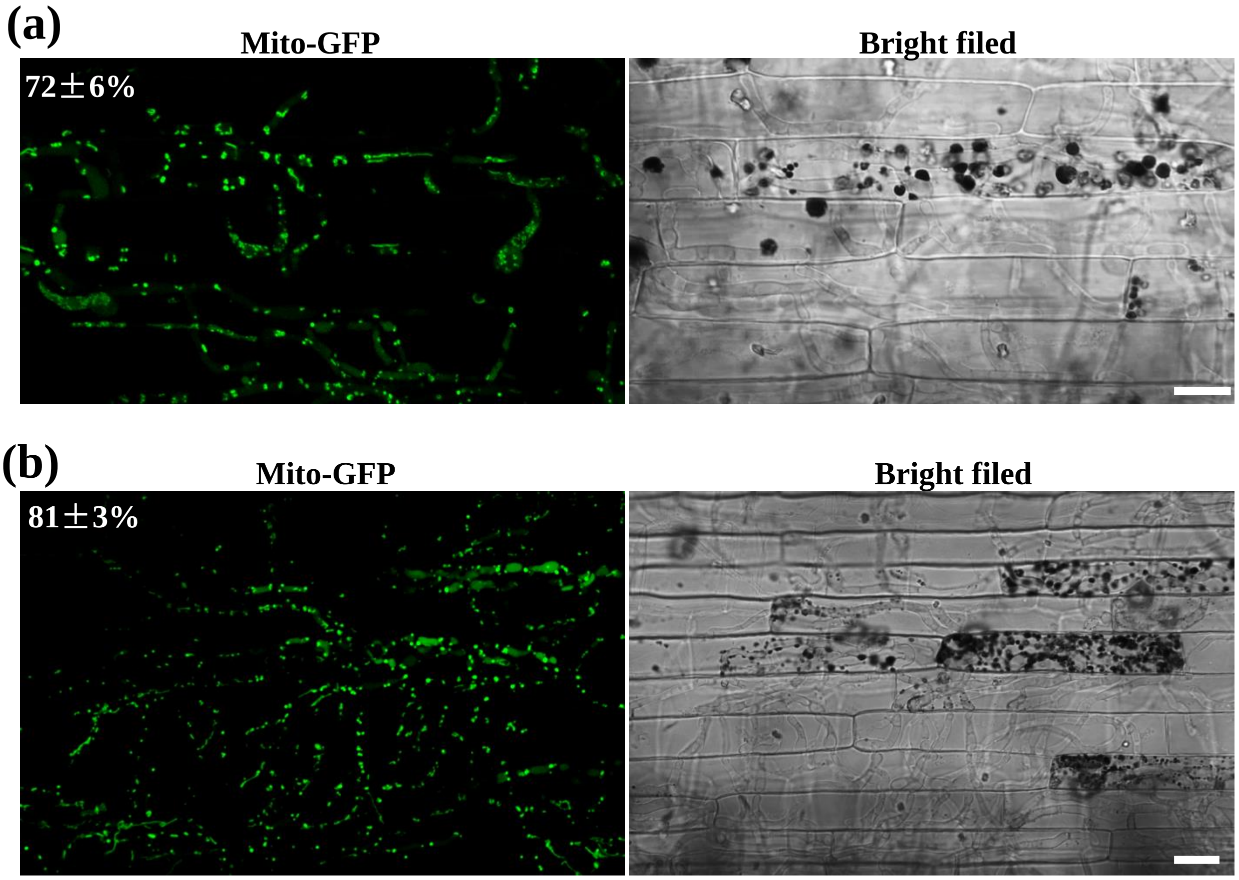


**Fig. S8** $NH_4NO_3$ or GR24 treatment did not change the mitochondrial fragmentation during infection by *M. oryzae*. Confocal microscopy image of the *Mito-GFP* strain in rice sheath cells in present of $NH_4NO_3$ (a) and GR24 (b). A conidial suspension of the *Mito-GFP* strain was inoculated on the rice sheath for 24 h before 1g/L $NH_4NO_3$ or 20 μM GR24 was added. Confocal microscopy was carried out at 48 hpi. Scale bar: 10 μm. The numbers on the pictures represent filamentous/total mitochondria (mean ± SD) from three independent experiments. Sample size is more than 50 appressoria penetration sites per analysis.

### Fig. S9

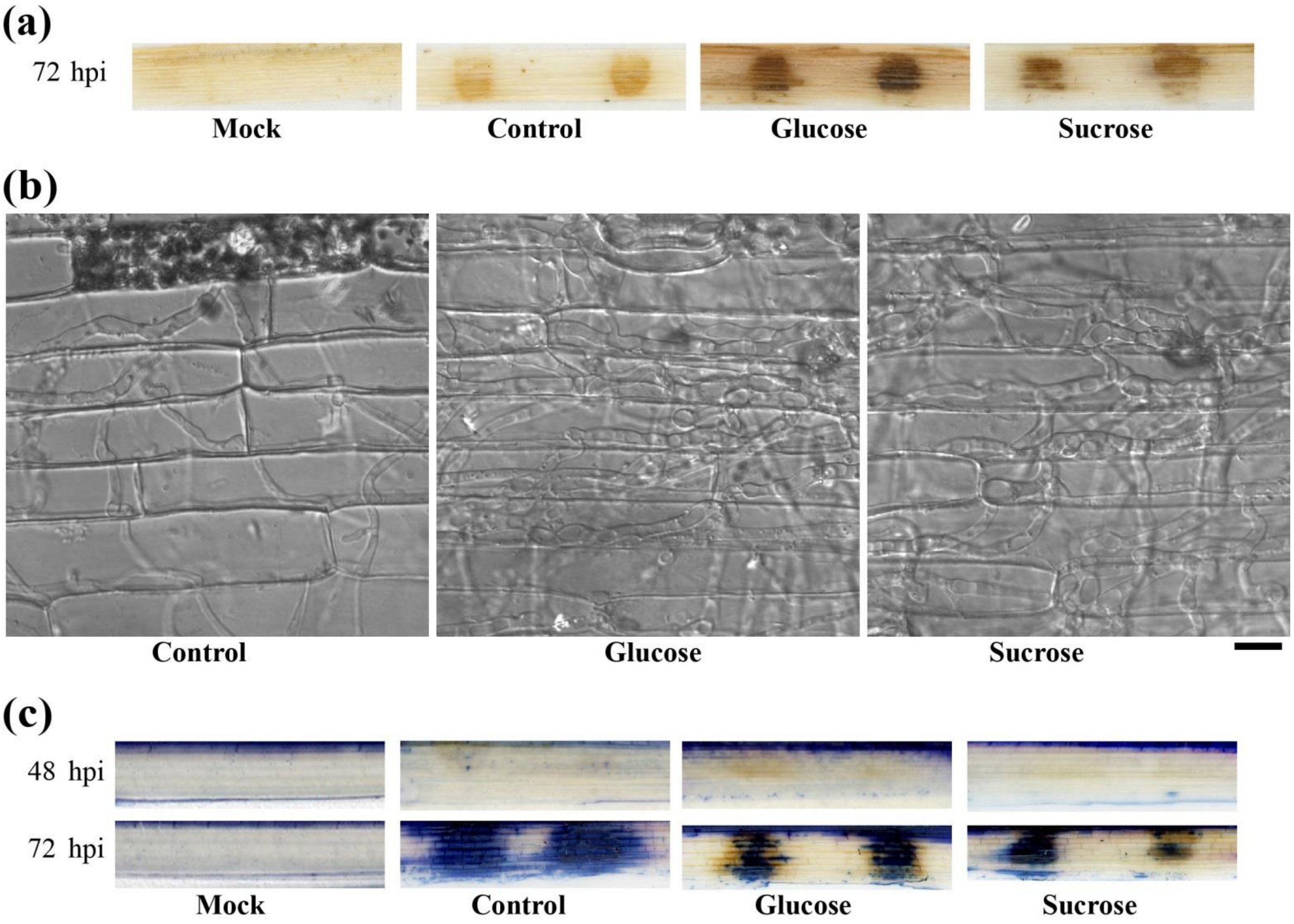


**Fig. S9** Addition of glucose or sucrose promotes spread of invasive hyphae and decreases the cell death in rice during early infection by *M. oryzae*. (a) The formation of blast disease lesions by the *Mito-GFP* strain in the presence of water (Mock), 8 mg/mL sucrose, or 50 mg/mL glucose. (b) The invasive hyphal growth of the *Mito-GFP* strain at 72 hpi with or without additional carbon source. In control (without glucose or sucrose), the collapsed of the first invaded cell was evident, whereas this did not occur in the presence of sucrose or glucose at 72 hpi. Bar = 10 μm. (c) Trypan blue staining highlights the cell death in infected rice sheath. Mock, no conidia; Control, conidia with $H_2O$ were inoculated on the rice sheath; Glucose or Sucrose, the water from conidial suspension was removed and supplement with glucose or sucrose at 24 hpi. Trypan blue staining was performed by immersing the cleared rice sheaths in the staining solution (0.01 % trypan blue in lactophenol) at room temperature for at least 4 h. The stained rice sheaths were stored in 60 % glycerol until microscopic observation.
